# Ageing leads to nonspecific antimicrobial peptide responses in *Drosophila melanogaster*

**DOI:** 10.1101/2022.06.25.497570

**Authors:** Biswajit Shit, Arun Prakash, Saubhik Sarkar, Pedro F. Vale, Imroze Khan

## Abstract

Evolutionary theory predicts a late-life decline in the force of natural selection, possibly leading to late-life deregulations of the immune system. A potential outcome of such immune-deregulation is the inability to produce specific immunity against target pathogens. We tested this possibility by infecting multiple *Drosophila melanogaster* lines (with bacterial pathogens) across age-groups, where either individual or different combinations of Imd- and Toll-inducible antimicrobial peptides (AMPs) were deleted using CRISPR gene editing. We show a high degree of non-redundancy and pathogen-specificity of AMPs in young flies: in some cases, even a single AMP could confer complete resistance. In contrast, ageing led to a complete loss of such specificity, warranting the action of multiple AMPs across Imd- and Toll-pathways during infections. Moreover, use of diverse AMPs either had no survival benefits, or even accompanied survival costs post-infection. These features were also sexually dimorphic: females expressed a larger repertoire of AMPs than males, but extracted equivalent survival benefits. Finally, age-specific expansion of the AMP-pool was associated with downregulation of negative-regulators of the Imd-pathway and a potential damage to renal function, as features of poorly-regulated immunity, Overall, we could establish ageing as an important driver of nonspecific AMP responses, across sexes and bacterial infections.

## INTRODUCTION

Ageing often leads to physiological senescence, including immune senescence, characterised by exaggerated and over-reactive pro-inflammatory responses (Stout-Delgado et al., 2009; Khan et al., 2017). In several insects (e.g. fruit flies and flour beetles), older individuals show increased expression of antimicrobial peptides (AMPs) (Zerofsky et al., 2005); higher haemolymph antibacterial activity or phenoloxidase response (PO) after infection, without any significant survival benefits (Khan et al., 2016). Instead, increased immunity often induces lethal immunopathological damage in older individuals, increasing their mortality rate (Khan et al., 2017; Badinloo et al., 2018). Similar effects are also reported in vertebrates, where an increase in chronic inflammatory state with age leads to maladaptive impacts of the innate immune system (Shaw et al., 2013). For example, older mice die faster owing to an elevated level of interleukin-17 and neutrophil activation, causing hepatocyte necrosis (Stout-Delgado et al., 2009). By and large, older individuals are thus more likely to experience the detrimental effects of overactive immunity in both invertebrate and vertebrate species.

Age-specific hyper-activation of immunity is consistent with the evolutionary theory of ageing which predicts a progressive decline in the force of natural selection with age (Williams, 1957; Hamilton, 1966)— natural selection that optimizes organismal physiology for development and reproduction early in life, can become too weak to effectively regulate the late-life performance in older individuals (Maklakov and Chapman, 2019). For example, poor regulatory mechanisms in several evolutionarily conserved signalling pathways such as insulin/ insulin-like growth factor signalling can result in suboptimal levels of gene expression in late life, with myriad negative health effects (Kenyon, 2010; Flatt and Partridge, 2018; Carlsson et al., 2021). Such changes in conserved signalling pathways might also interfere with the optimal induction and regulation of costly immune pathways in aged individuals. Several experiments on age-specific changes in the expression of negative regulators of immunity support this hypothesis (Neves and Sousa-Victor, 2020): e.g., reduced expression of anti-inflammatory cytokine interleukin-10 not only causes over-activation of cytotoxic inflammatory pathways in older mice, but also promotes their muscular, cardiovascular and metabolic dysfunction (Mohanty et al., 2015). In older mice and humans, a rapid age-specific decline of another immunomodulatory molecule, MANF, increases the levels of pro-inflammatory cytokines and activated macrophages (Mohanty et al., 2015; Neves et al., 2016). These changes in immunity and fitness effects are thus an outcome of age-related malfunctioning of regulatory units of immune pathways.

A further potential manifestation of such a deregulated ageing immune system is the progressive loss of specificity to pathogens. Younger individuals can optimise their immune responses by acting selectively on pathogens with a limited set of immune effectors (Moret, 2003, Hanson et al. 2019). In contrast, older individuals, owing to their poorly regulated immunity, might show nonspecific activation of higher number of immune effectors against an equivalent dose of antigenic exposure. An extended immune repertoire can also collectively increase the cytotoxicity of immune responses, elevating the risk of morbidity and mortality with ageing (Khan et al., 2017; Badinloo et al., 2018). Indeed, prior experiments with older mice showed that pathways leading to increased production of antigen non-specific antibodies can enhance the risk of autoimmune responses with no improvement in pathogen clearance ability or survival (Bruce et al., 2009). However, experiments measuring the functional expansion of the available immune repertoire with ageing and their role in overall infection outcome is currently missing.

In the present work, we tested the impact of ageing on specific interactions between immune effectors and bacterial infections, using multiple *D. melanogaster* lines where different combinations of AMPs from the Imd and Toll pathways were knocked out by CRISPR/Cas9 gene editing (Hanson et al., 2019). We targeted AMPs as they have been recently shown to possess a high degree of non-redundancy, non-interchangeability and specificity against a range of pathogens in young flies (Hanson et al., 2019). Only a small subset of the total AMP repertoire provides the most effective protection against specific pathogens so that in some cases, even a single AMP is sufficient to control the growth of specific pathogens: e.g., Imd-pathway responsive AMP *Diptericins* (or *Drosocin*) against *Providencia rettgeri* (or *Enterobacter cloacae*) infection. Such specificity of AMP responses might also indicate potentially higher adaptive values associated with using fewer immune effectors in young individuals, thereby, avoiding the net fitness costs of general immune activation (Moret, 2003). Indeed, earlier experiments suggest that toxic levels of AMP expression, due to suppression of negative immune regulators of Imd-pathway (or increased transcriptional activation of its positive regulators) in young flies, can lead to reduced lifespan or extensive neurodegeneration causing faster ageing (Kounatidis et al., 2017). We speculate that the general loss of regulation in an ageing immune system might also accompany loss of such controlled specific AMP actions, deploying more AMPs to counter equivalent infection levels, but without any added survival benefits.

In addition, the age-specific role of AMPs can be sex-specific with a strong sex-by-age interactions (Belmonte et al., 2020). For instance, overexpression of relish or Toll-responsive defensin can reduce male lifespan more than that of females in *Drosophila* (Badinloo et al., 2018). Also, previous studies with flies infected with *P. rettgeri* indicated *Drosophila* males had higher *Diptericin* expression (Duneau et al., 2017). A relatively higher expression of *Diptericin* transcripts in males is perhaps needed to support its exclusive role against *P. rettgeri* infection, whereas low expression in females opens up possibilities where *Diptericin* is either dispensable or requires compensatory actions of other AMPs. However, there are no direct experiments to test these possibilities of sex-specific expansion of AMP use.

## MATERIALS AND METHODS

### I. Fly strains and maintenance

To test the role of ageing on AMP-driven specific immunity, we used multiple *Drosophila melanogaster* lines where different combinations of multiple and individual AMPs were knocked out mostly using the CRISPR/Cas9 gene editing or homologous recombination (details described in Hanson et al., 2019; also see Fig. S1). We used null mutants for 10 of the 14 known *Drosophila* AMPs that are expressed upon systemic infection. These include mutations from six single gene families including *Defensin* (*Def^SK3^*), *Attacin C* (*AttC^Mi^*), *Attacin D* (*AttD^SK3^*), *Drosocin* (*Dro^SK4^*) *Metchnikowin* (*Mtk^R1^*) and *Drosomycin* (*Drs^R1^*) loci and two small deletion removing *Diptericins DptA* and *DptB (Dpt^SK1^)*, or the gene cluster containing *Drosocin* and *Attacins AttA* and *AttB* (*Dro*-*AttAB^SK2^*). The iso-*w^1118^* (DrosDel isogenic) wild-type was used as the genetic background for mutant isogenization (see Ferreira et al., 2014; Hanson et al., 2019). We also used ‘Δ*AMPs’* flies where independent mutations were recombined into a background lacking 10 inducible AMPs. However, we note that the impact of *ΔAMPs* could be due to AMPs having specific effects or combinatorial action of multiple co-expressed AMPs. To tease apart these effects, we also included various combined mutants where different groups of AMPs were deleted based on the pathways that they are controlled by: (1) Group B - flies lacking AMPs such as *Drosocin*, *Diptericins* and *Attacins* (*AttC^Mi^*; *AttD^SK1^*; *Dro^SK4^*; *Dro*-*AttAB^SK2^*) (exclusively regulated by Imd-pathway) (2) Group C - flies lacking the two Toll-regulated antifungal peptide genes *Metchnikowin* and *Drosomycin* (*Mtk^R1^*; *Drs^R1^*) (mostly regulated by Toll-pathway). We also referred to flies with single mutations lacking *Defensin* (*Def^Sk3^*) (co-regulated by Imd- and Toll-pathway) as group A. Finally, we also included fly line where group-A, B and C mutants were combined to generate flies lacking AMPs either from groups A and B (AB), or A and C (AC), or B and C (BC).

We maintained all fly stocks and experimental individuals on a standard cornmeal diet also known as Lewis medium (Siva-Jothy et al., 2018) at a constant temperature of 25°C on a 12 : 12 hour light: dark cycle at 60% humidity. To generate the experimental flies, we reared flies at a larval density of ~70 eggs/ 6ml food. We collected adult males and females as virgins and held at a density of 25 flies/sex/food vial for the experiment described below. Female iso-*w^1118^* flies undergo reproductive senescence within 25 days post-eclosion (Reproductive output measured for 18-hours; Mean ± SE: 3-day-old= 6.75 ± 0.77 vs 24-day-old= 3.17 ± 0.53, P<0.001). Hence, in our experiments, we used 3 and 25-day-old individuals (post-eclosion) as ‘young’ and ‘old’ adults, respectively. We transferred the adults to fresh food vials every 3 days, during the entire experimental window. By screening the single mutants, along with combined genotypes, we were able to compare the changes in specific immunity as a function of possible interactions between AMPs of different groups’ vs function of individual AMPs with ageing.

### II. Infection protocol and the assay for post-infection survival

For all the infection, we either used Gram-negative bacteria *Providencia rettgeri* or *Pseudomonas entomophila.* Both are natural pathogens of *Drosophila* that activate the IMD pathway (Myllymäki et al., 2014) and could impose significant mortality (Galac and Lazzaro, 2011; Dieppois et al., 2015). To quantify post-infection survivorship, we infected flies (septic injury method) in the thorax region with a 0.1 mm minutien pin (Fine Science Tools) dipped into a bacterial suspension made from 5 mL overnight culture (optical density of 0.95, measured at 600 nm) of either *Providencia rettgeri* or *Pseudomonas entomophila* adjusted to OD of 0.1 and 0.05 respectively (See SI methods for details). In total, we infected 160-280 flies/sex/infection treatment/bacterial pathogen/age-group/fly genotypes and then held them in food vials in a group 20 individuals (For each treatment, sex, age-group, pathogen type, we thus had 8-14 replicate food vials). We carried out sham infection with a pin dipped in sterile phosphate buffer solution (1X PBS).

We then recorded their survival every 4-hours (±2) for 5 days. Due to logistical challenges of handling a large number of flies, we infected each sex and age-groups with *P. rettgeri* (or *P. entomophila*) separately in multiple batches, where they were handled as — (i) Groups AB, BC, AC; (ii) Group-A, B & C; (iii) Imd-responsive and (iv) Toll-responsive single mutants for *P. rettgeri*; or (i) Groups AB, BC, AC, A, B & C; (ii) Imd-responsive and (iii) Toll-responsive single mutants for *P. entomophila*. Every time, we also assayed iso-*w^1118^* flies as a control to facilitate a meaningful comparison across different batches. Therefore, although sexes and age-groups for each mutant were not directly comparable, their relative effects with respect to control iso-*w^1118^*were estimated across sexes, age-groups and pathogen types. Note that we compared each mutant separately with iso-*w^1118^* flies, since we only wanted to capture their changes in infection susceptibility relative to control flies. For each batch of flies, across pathogen types, sexes and age-groups, we analysed the survival data with a mixed effects Cox model, using the R package ‘coxme’ (Therneau, 2015). We specified the model as: survival ~ fly lines (individual AMP mutant lines vs iso-*w^1118^*) + (1|food vials), with fly lines as a fixed effect and replicate food vials as a random effect. Since none of the fly lines had any mortality after sham-infection, we were able to quantify the susceptibility of each infected mutant lines (AMP knockouts) with respect to control flies (iso-*w^1118^* group) as the estimated hazard ratio of infected AMP mutants versus control flies (hazard ratio = rate of deaths occurring in infected AMP mutants /rate of deaths occurring in iso-*w^1118^* group). A hazard ratio significantly greater than one indicated a higher risk of mortality in the AMP mutant individuals.

Note that the above experimental design allowed us to repeat the assay for post-infection survival of young and old iso-*w^1118^* flies infected with *P. rettgeri* (or *P. entomophila*) in 4 (or 3) independently replicated experiments. We thus estimated the effects of ageing on their post-infection survival, using a mixed effects Cox model specified as: survival ~ age + (1|food vials), with age as a fixed effect, and food vials as random effects.

### III. Assay for bacterial clearance

Mortality of control flies (iso-*w^1118^*) injected with the experimental infection dose began around 24-hours and 20-hours after infection with *P. rettgeri* and *P. entomophila* respectively (Fig. S2A). We therefore used these time-points to estimate the bacterial load across the age-groups as a measure of the pathogen clearance ability across AMPs (see SI methods for detailed protocol). We homogenized flies in a group of 6 in the sterile PBS (n= 8-15 replicate groups/sex/treatment/age-group/fly lines), followed by plating them on Luria agar. Due to logistical challenges with large number of experimental flies, we handled each sex, age-group and pathogen type separately and in multiple batches as described above.

Also, similar to post-infection survival data, we were only interested in comparing the changes in bacterial load for each mutant line relative to control iso-*w^1118^*flies across experimental groups. We thus analysed the bacterial load data of each mutant genotype with iso-*w^1118^* flies separately across age-groups, sexes and pathogen types. Since residuals of bacterial load data were non-normally distributed (confirmed using Shapiro-Wilks’s test), we log-transformed the data, but residuals were still non-normally distributed. Subsequently, we analysed the log-transformed data, using a generalised linear model best fitted to gamma distribution, with fly lines (i.e., control iso-*w^1118^* line vs individual AMP knockout line) as a fixed effect.

### IV. Assay for the Malpighian tubule activity, as a proxy for immunopathological damage

Malpighian tubules (MTs), the fluid-transporting excretory epithelium in all insects, are prone to increased immunopathology following an immune activation due to their position in the body and the fact that they cannot be protected with an impermeable membrane due to their functional requirement (Dow et al., 1994; Khan et al., 2017). Previous experiments have shown that risk of such immunopathological damage can increase further with ageing in mealworm beetle *Tenebrio molitor* (Khan et al., 2017). It is possible that nonspecific AMP responses with ageing in *Drosophila* was also associated with increased immunopathological damage to MTs. We thus estimated the fluid transporting capacity of functional MTs dissected from experimental females at 4-hours after immune challenge with 0.1 OD *P. rettgeri* (n=12-20 females/ infection treatment/age-group), using a modified ‘oil drop’ technique as outlined in previous studies (Dow et al. 1994; Li et al., 2020) (also see SI methods).

This method provides a functional estimate of their physiological capacity by assaying the ability to transport saline across the active cell wall into the tubule lumen. The volume of the secreted saline droplet is negatively correlated with the level of immunopathological damage to MTs. Since we collected the flies across the age-groups on different days, we analysed the MT activity data as a function of infection status for each age-group separately, using a generalized linear mixed model best fitted to a quasibinomial distribution.

### V. Gene expression assay

Finally, we note that transcription of negative regulators of Imd-pathway such as *pirk* and *caudal* are important to ensure an appropriate level of immune response following infection with gram-negative bacterial pathogens, thereby avoiding the immunopathological effects (Lee and Ferrandon, 2011; Kleino and Silverman, 2014). While *pirk* interferes with the interaction of *PGRP-LC* and *-LE* with the molecule Imd to limit the activation of the Imd pathway, *caudal* downregulates the expression of AMPs (Lee and Ferrandon, 2011). To examine whether non-specific expansion of AMP repertoire was associated with the lower expression of these negative regulators, we estimated their relative expression level in both young and old iso-*w^1118^*individuals infected with *P. rettgeri* at 24 hours post-infection, by using qPCR (as outlined in Prakash et al., 2021) (n= Total 15-21 flies in a group of 3 homogenized in Trizol reagent/ Infection treatment/ age-group and sex-combination).

In addition, we also estimated the expression of the Imd-pathway NF-κB transcription factor *Relish* and peptidoglycan recognition protein - *PGRP-LC*, both act as positive regulators of Imd-pathway (Lemaitre and Hoffmann, 2007; Myllymäki et al., 2014) and hence, can serve as a proxy for overactivated Imd pathway and higher AMP expression in older flies (Badinloo et al., 2018) (also see SI methods). We analysed the gene expression data using ANOVA (see SI methods section-iv for details).

## RESULTS

We began our observation with high mortality and increased bacterial load in AMP-deficient flies (Δ*AMPs*) infected with *P. rettgeri*, regardless of their sex and age (Fig. 1A, 1B, 1E, 1F; Table S2, S3), suggesting that AMPs are critically important to prevent pathogen growth, thereby increasing the post-infection survival costs (Fig. 1A, 1B, 1E, 1F; Fig. S3, S4; Table S2, S3). Older iso-*w^1118^* control females infected with *P. rettgeri* also showed higher mortality (Fig. S2A, S2B; Table S4A) and increased bacterial load (Fig. S2C, Table S4B) than their younger counterparts, suggesting negative effects of ageing on fitness and pathogen clearance ability. By contrast, both young and old males had similar post-infection mortality rate (Fig. S2A, S2B; Table S4A) with comparable bacterial load (Fig. S2C; Table S4B), indicating that ageing did not impact male’s ability to survive post-infection and clear pathogens, at least at the infection dose used in our experiments. Nevertheless, in the subsequent assays, these results from male flies infected with *P. rettgeri* enabled us to directly compare the relative effects of deleting different Imd- vs Toll-responsive AMPs across age-groups (against a common baseline).

**Figure 1.**
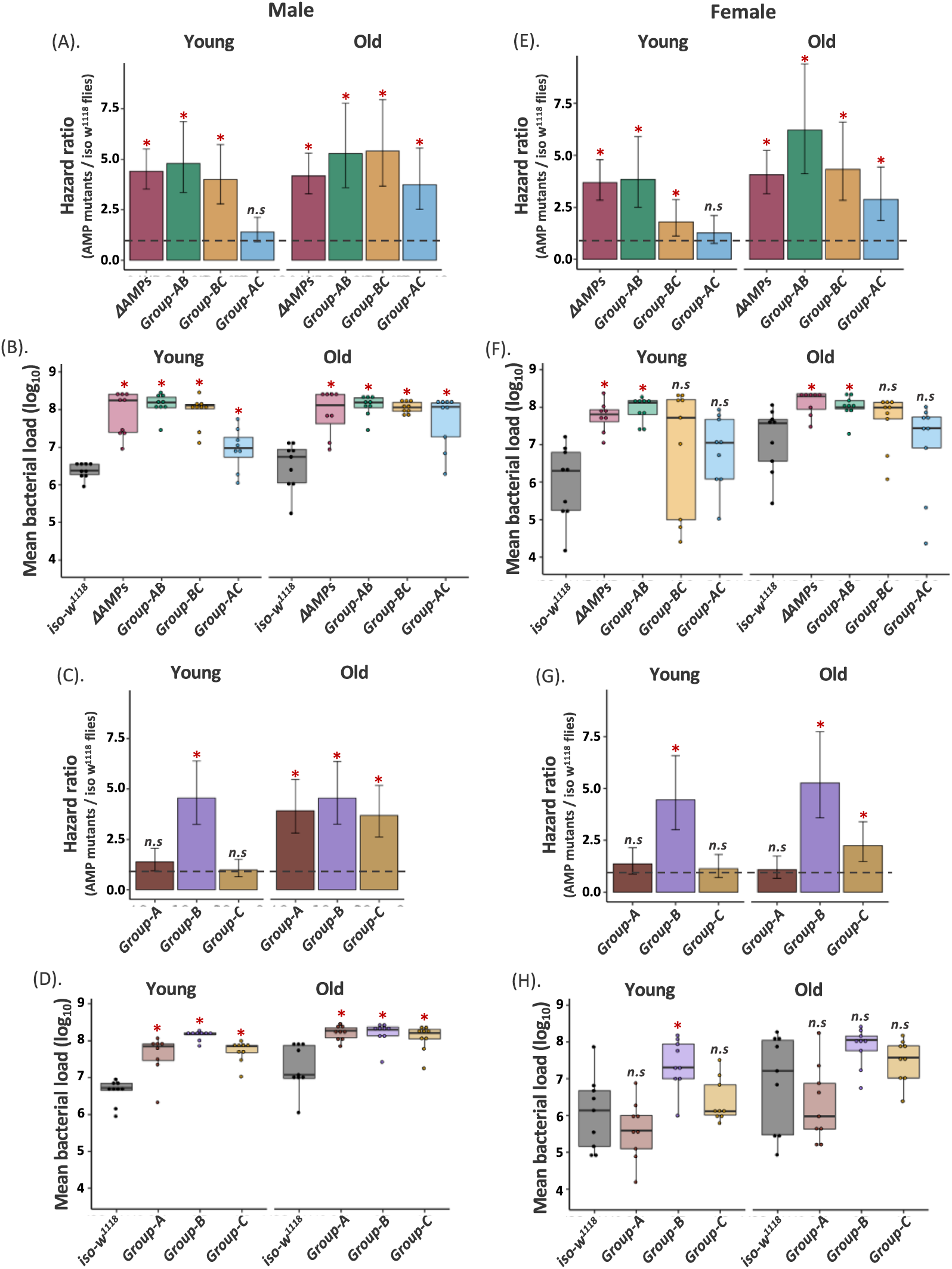
Infection with *Providencia rettgeri* in multiple AMP-knockouts. The estimated hazard ratios calculated from survival curves (160-180 flies/sex/infection treatment/age-group/fly line; see Fig. S3, S4) and bacterial load (n= 8-9 replicate groups/sex/treatment/age-group/fly line) measured at 24-hours after *P. rettgeri* infection across sexes and age-groups. Hazard ratios for double combination of AMP-knockouts (i.e., group-AB, BC, & AC; see Fig. S1 for details about the fly lines) in males **(A)** and females **(E)**. Bacterial loads for double combination of AMP-knockouts in males **(B)** and females **(F)**. Hazard ratios for single combination of AMP Knockouts (e.g., group-A, B & C) in males **(C)** and females **(G).** Bacterial load for single combination of compound of AMP-knockouts in males **(D)** and females **(H)**. In panels A, C, E, G, hazard ratios significantly greater than 1 (hazard ratio =1; shown as horizontal dashed grey lines), indicated by asterisk (*), suggests higher infection susceptibility of mutant flies than the iso-*w^1118^* control flies. In panels B, D, F, H, each data point represents the bacterial load of flies pooled in a group of 6. Mutant fly lines that had significantly different bacterial load from wild-type iso-*w^1118^* are indicated by asterisks. ns = not significant. Group A-flies lacking *Defensin*; Group B - flies lacking AMPs such as *Drosocin*, *Diptericins* and *Attacins*; Group C - flies lacking *Metchnikowin* and *Drosomycin*; Group-A, B and C mutants were combined to generate flies lacking AMPs either from groups A and B (AB), or A and C (AC), or B and C (BC).

### I. Ageing leads to an expansion of the required AMP repertoire against *P. rettgeri* infection

To gain a broad understanding of how AMP specificity changes with age, we first tested mutants lacking different groups of AMPs either from Imd- (e.g., group B) or Toll-pathways (e.g., group C) (pathway-specific), or combined mutants lacking pathway-specific mutants in different combinations (e.g., group AB, BC or AC) (See Fig. S1 for description of mutants). As reported in a previous study by Hanson and co-workers (Hanson et al., 2019), young males lacking group-AB and -BC AMPs were highly susceptible to *P. rettgeri* infection (Fig. 1A; Table S2), and this was generally associated with 10-100-fold increased bacterial loads in these mutants relative to the iso-*w^1118^* control (Fig. 1B; Table S3). Subsequent assays with pathway-specific (i.e., Imd- or Toll-pathway) AMP combinations (group A, B or C) confirmed that such effects were primarily driven by Imd-regulated group-B AMPs that were shared between both AB and BC combinations (Fig. 1C; Fig. S3; Table S2), and equally driven by increased bacterial load (Fig. 1D; Table S3). We found a comparable pattern in young females as well, except that flies lacking group BC combinations of AMPs were not negatively affected by infection (Fig. 1E, 1F, 1G, 1H; Fig. S4; Table S2, S3).

In contrast to young flies, most of the pathway-specific or combined mutants became highly susceptible to *P. rettgeri* infection with age, except females of group-A mutants flies lacking *Def*. This would suggest a possible sexually dimorphic effect of *Defensin* in *P. rettgeri* infection, which appear to be important for males, but not females (Fig. 1C, 1G; Fig. S3, S4; Table S2). Regardless of this slight variation across sexes, our results clearly demonstrated that only having functional Imd-regulated group-B AMPs was not sufficient to protect older flies against *P. rettgeri* infection. Also, these results indicated that a single AMP *Dpt*-driven protection against *P. rettgeri* infection, as suggested by Hanson et al. (2019), may not be applicable to older flies (also see Unckless et al., 2016). High susceptibility of older mutants lacking group A or C AMPs (Fig. 1C, 1G; Table S2) and increased bacterial growth (Fig. 1D, 1H; Table S3) therein clearly indicated that other AMPs responsive to Gram-positive bacteria (e.g., *Def*) or fungal pathogens (e.g., *Mtk*, *Drs*) might be needed as well.

### II. *Dpt*-specificity against *P. rettgeri* infection is sex-specific and disappears with age

Next, we decided to test the role of individual AMPs deleted in the pathway-specific or compound mutants across age-groups and sexes. Interestingly, *Dpt* provided complete protection against *P. rettgeri* only in young males, but not in females or older males (Fig. 2A, 2E; Table S5). This was verified by using fly lines where *DptA* and *DptB* are introduced on an AMP-deficient background (Δ*AMPs^+Dpt^*). *Dpt* reintroduction could fully restore survival as that of wild-type flies only in young males, and this was associated with a decrease in CFUs compared to the *Dpt* deletion mutant (Fig. 2B, Table S6). However, reintroduction of functional *DptA* and *DptB (*Δ*AMPs^+Dpt^*) in young or old females flies did not result in lower CFUs (Fig. 2F; Table S6) and these flies remained highly susceptible to *P. rettgeri* (Fig. 2E; Table S2). Young (or old) females also showed increased bacterial loads and associated higher susceptibility when other Imd-regulated group-B AMPs such as *AttC* and *Dro-Att* (or *Dro* and *Dro-Att* in old females) were deleted (Fig. 2E, 2F; Table S5, S6). Older females lacking *AttC* showed increased infection susceptibility as well, but did not have increased bacterial load (Fig. 2E, 2F; Table S5, S6). Older Δ*AMPs ^+Dpt^* males, on the other hand, could limit the bacterial burden as low as that of the control iso-*w^1118^* flies (Fig. 2B; Table S6), but still showed very high post-infection mortality (Fig. 2A; Table S5). These results from older males thus suggested that the ability to clear pathogens might not always translate into an improved ability to survive after infection (Fig. 2A, 2B; Table S5, S6).

**Figure. 2.**
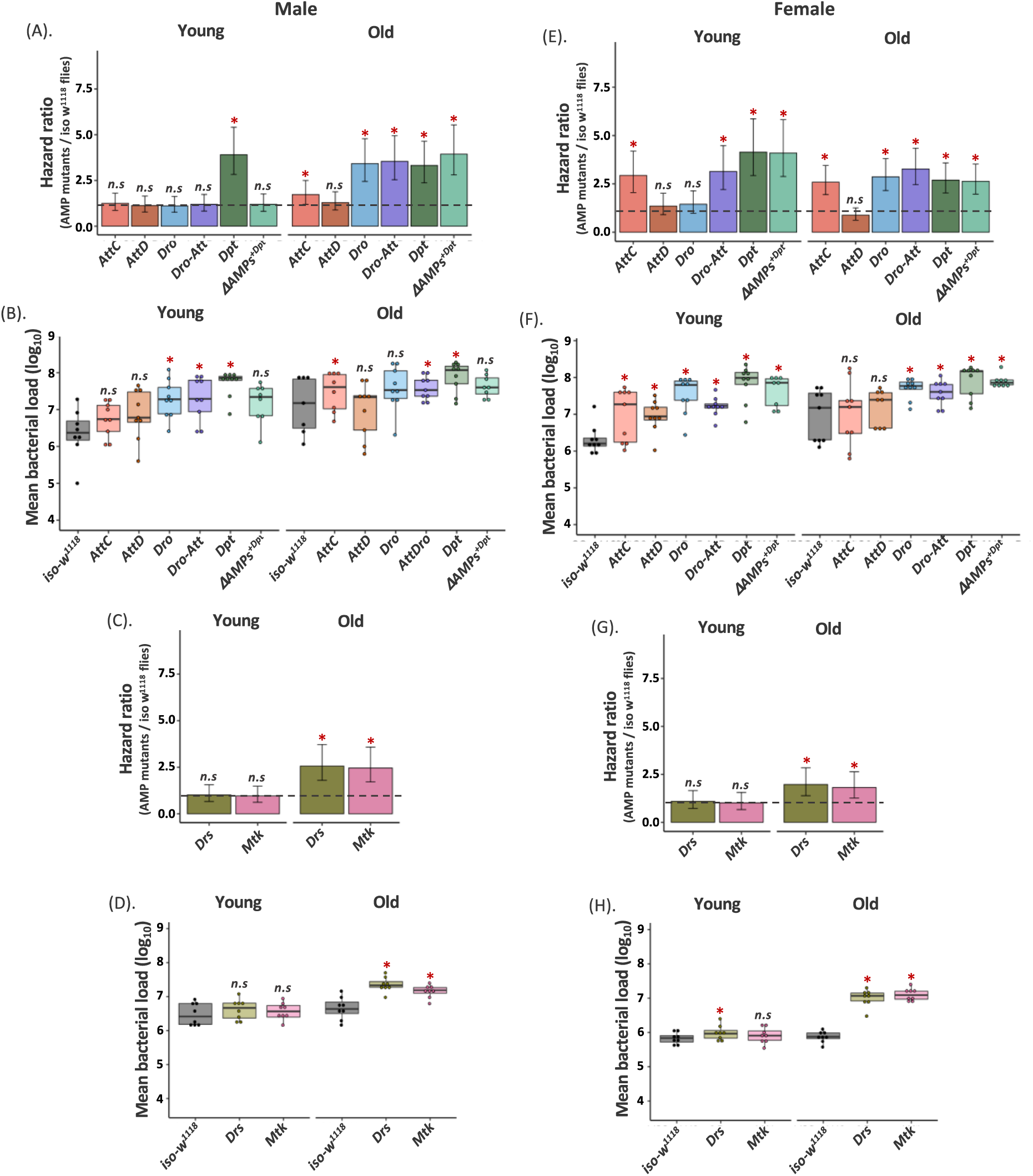
Infection with *Providencia rettgeri* in individual AMP-knockouts. The estimated hazard ratios calculated from survival curves (160-180 flies/sex/infection treatment/ age-group/fly line; see Fig. S3, S4) and bacterial load (n= 8-9 replicate groups/sex/treatment/age-group/fly line) measured at 24 hours after *P. rettgeri* infection across sexes and age-groups. Hazard ratios for Imd-responsive single AMP (e.g., *Dpt, AttC, AttD, Dro*; see Fig. S1 for details about the fly lines) and Att-Dro knockouts in males **(A)** and females **(E)**. Bacterial load of Imd-responsive single AMP and Att-Dro knockouts in males **(B)** and females **(F)**. Hazard ratios for Toll-responsive single AMP knockouts (e.g., *Drs & Mtk*) in males **(C)** and females **(G)**. Bacterial loads of Toll-responsive single AMP knockouts in males **(D)** and females **(H)** respectively. In panels A, C, E, G, hazard ratios significantly greater than 1 (hazard ratio =1; shown as horizontal dashed grey lines), indicated by asterisk (*), suggests higher infection susceptibility of mutant flies than the iso-*w^1118^* control flies. In panels B, D, F, H, each data point represents the bacterial load of flies pooled in a group of 6. Mutant fly lines that had significantly different bacterial load from wild-type iso-*w^1118^* are indicated by asterisks (*). ns = not significant.

Why did females always require AMPs other than *Dpt* after *P. rettgeri* infection? Although the mechanisms behind sex-specific expansion of AMP repertoire are unknown, a possible explanation is that females show inherently lower expression level of *Dpt* relative to males. Consequently, they may require the joint expression of other AMPs to complement the lower *Dpt* expression, thereby enhancing the protection against *P. rettgeri* infection. Indeed, a previous study has already demonstrated lower *Dpt* expression in iso-*w^1118^*females than males after *P. rettgeri* infection (see Duneau et al., 2017), although the causal link between reduced *Dpt* expression and proportional increase in the compensatory action of other AMPs is not yet experimentally validated.

Also, both males and females showed further extension to a Toll-responsive AMP repertoire with ageing. In addition to the role of *Def* (included in group-A) as described above in older males (but not in older females; compare Fig. 1D, 1H; Table S3), older flies of both sexes also showed increased microbe loads and increased mortality when Toll-regulated AMPs from such as *Drs* and *Mtk* were deleted (Fig. 2D, 2H; Table S6), raising a possibility of crosstalk between Toll and Imd immune-signalling pathways (Duneau et al., 2017; Nishide et al., 2019). Taken together, these results describe ageing as a major driver behind the loss of specificity of AMP responses.

Additionally, we also note that a few other mutations such as deletion of *Dro* and *Dro-Att*, which otherwise had no effects on the survival of *P. rettgeri*-infected young males, caused significant increase in the bacterial load (Fig. 2A, 2B; Table S5, S6). Together, these results not only underscored the multifaceted role of AMPs, but also provided functional resolution at the level of single AMPs such as *Dpt* which in addition to playing the canonical role in resisting the infection, also aided in withstanding the effects of increased pathogen growth, caused by the dysfunction of other AMPs (Fig. 2A, 2B; Table S5, S6).

### III. Expansion of the required AMP repertoire does not improve, and even reduces, survival in both older males and females infected with *P. entomophila*

To test if age-related loss of AMP specificity was specific to *P. rettgeri*, or also occurred with other infections, we investigated the AMP repertoire in young and old flies infected with the Gram-negative bacteria *P. entomophila*. Similarly, older flies required a larger repertoire of AMPs (Fig. 3A, 3C; Table S7) and yet, died faster than young flies (old vs young: 4-fold vs 2-fold; Fig. 3A, 3C). In contrast to younger flies, where only group-B, -AB and -BC mutants were susceptible to *P. entomophila* infection, all the other pathway-specific or combined mutants of older males and females were also highly sensitive to infection (Fig. 3A, 3C; Table S7). However, further experiments with single AMP mutants revealed that the antibacterial protection in both young males and females was still limited only to the exclusively Imd-regulated group-B AMPs, where several of them individually caused significant increase in microbe loads and reduction in post-infection survival (Fig. 4A, 4B, 4E, 4F; Table S9, S10). In contrast, older flies also needed additional action of Toll-regulated AMP *Drs* (Fig. 4C, 4G; Table S9), though it is striking that that increased mortality was not associated with increased microbe loads relative to iso-*w^1118^* in this case (Fig. 4D, 4H; Table S10). Overall, this is comparable to *P. rettgeri* infection where potential crosstalk between Toll & Imd immune-signalling pathways has already been implicated with ageing (Fig. 2; Table S5, S6). Also, the broad similarity between age-specific expansion and cross-reactivity of AMP repertoire against two different pathogens indicated the possibility where non-specificity can indeed be a generalised feature of an ageing immunity. Moreover, the increased mortality in older flies infected with *P. entomophila*, despite involving a higher number of AMPs, was perhaps an indication of their exacerbated cytotoxic effects with age (Badinloo et al., 2018).

**Figure 3.**
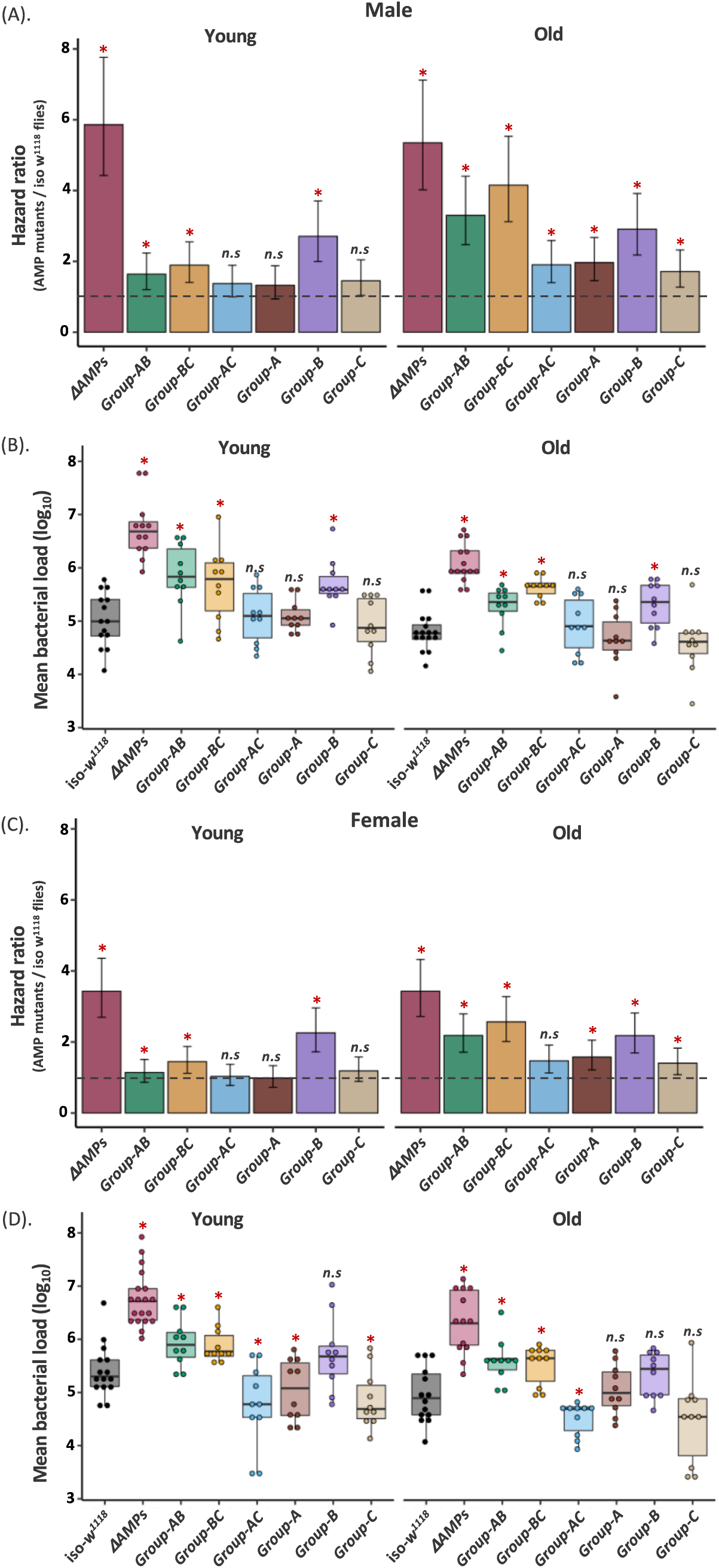
Infection with *Pseudomonas entomophila* in multiple AMP-knockouts. The estimated hazard ratios calculated from survival curves (180-280 flies/treatment/age-groups/sex/fly line; see SI Fig. S5, S6) and bacterial load (n= 9-15 replicate groups/sex/treatment/age-group/fly line) measured at 20-hours after *P. entomophila* infection across sexes and age-groups. Hazard ratios for double (i.e., group-AB, BC, & AC) and single combination (i.e., group-A, B, C) of AMP-knockouts in males **(A)** and females **(C)**. Bacterial loads for double and single combination of AMP-knockouts in males **(B)** and females **(D)**. In panels A & C hazard ratios significantly greater than 1 (hazard ratio =1; shown as horizontal dashed grey lines), indicated by asterisk (*), suggests higher infection susceptibility of mutant flies than the iso-*w^1118^* control flies. In panels B & D each data point represents the bacterial load of flies pooled in a group of 6. Mutant fly lines that had significantly different bacterial load from wild-type iso-*w^1118^* are indicated by asterisks. ns = not significant. See Fig 1 or the main text for the description of different fly groups.

**Figure 4.**
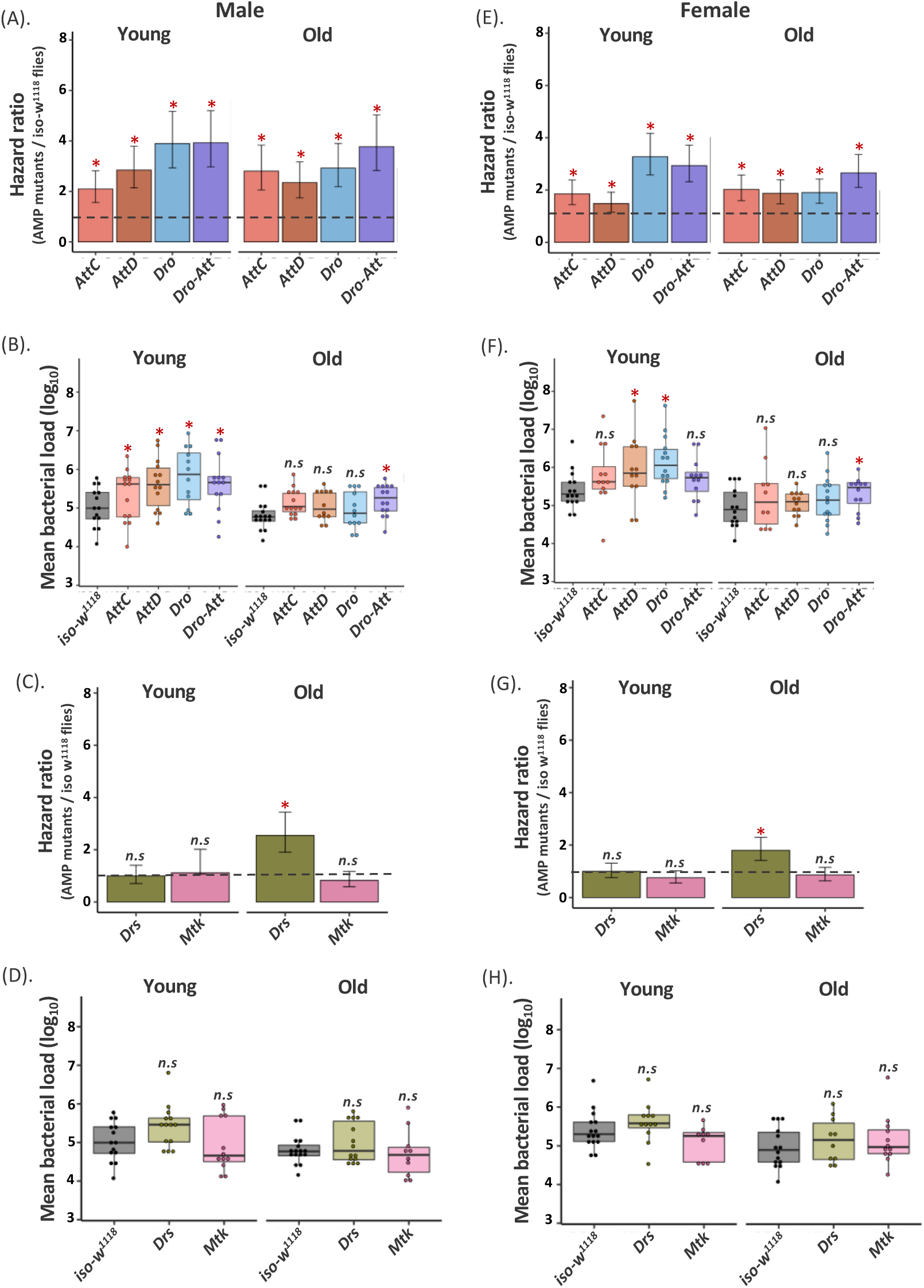
Infection with *Pseudomonas entomophila* in individual AMP-knockouts. The estimated hazard ratios calculated from survival curves (180-280 flies/sex/infection treatment/ age-group/fly line; see SI Fig. S5, S6) and bacterial load (n= 9-15 replicate groups/sex/treatment/age-group/fly line) measured at 20 hours after *P. entomophila* infection across sexes and age-groups. Hazard ratios for Imd-responsive single AMP (e.g., *AttC, AttD, Dro*; see Fig. S1 for details about the fly lines) and *Att-Dro* knockouts in males **(A)** and females **(E)**. Bacterial load of Imd-responsive single AMP and *Att-Dro* knockouts in males **(B)** and females **(F)**. Hazard ratios for Toll-responsive single AMP knockouts (e.g., *Drs & Mtk*) in males **(C)** and females **(G)**. Bacterial loads of Toll-responsive single AMP knockouts in males **(D)** and females **(H)** respectively. In panels A, C, E, G, hazard ratios significantly greater than 1 (hazard ratio =1; shown as horizontal dashed grey lines), indicated by asterisk (*), suggests higher infection susceptibility of mutant flies than the iso-*w^1118^* control flies. In panels B, D, F, H, each data point represents the bacterial load of flies pooled in a group of 6. Mutant fly lines that had significantly different bacterial load from wild-type iso-*w^1118^*are indicated by asterisks (*). ns = not significant.

### IV. Ageing-induced expansion of the required AMP repertoire was associated with downregulation of negative immune regulators and a trend of reduced renal purging post-infection

The expansion of the AMP repertoire in older flies could reflect a compensatory action to balance the lower per capita efficiency of their individual AMPs. This would enable flies to maintain an equivalent post-infection survival as that of younger flies against similar infection dose (e.g., old vs young males infected with *P. rettgeri*; Fig. 1A; Table S2). However, any benefits of recruiting multiple AMPs, may have been outweighed by the costs of expressing them (suggested in Badinloo et al., 2018) as higher immune activity, in general, accelerates the ageing process by imposing immunopathological damage to vital organs such as Malpighian tubules (MTs) (Khan et al., 2017). We expected that expansion of the AMP repertoire might have similar consequences in our experimental older flies as well. This is closely reflected by our results where *P. rettgeri* infection significantly reduced renal function in older females, measured as MT secretion (Fig. 5A; Table S11). Since functional MTs are needed to purge excessive ROS produced during immune responses as a physiological adaptation to prevent tissue damage in *Drosophila* (Li et al., 2020), reduced MT activity might not only exacerbates the effects of pathogenic infection, but can also causes late-life costs (Khan et al., 2017).

**Figure 5.**
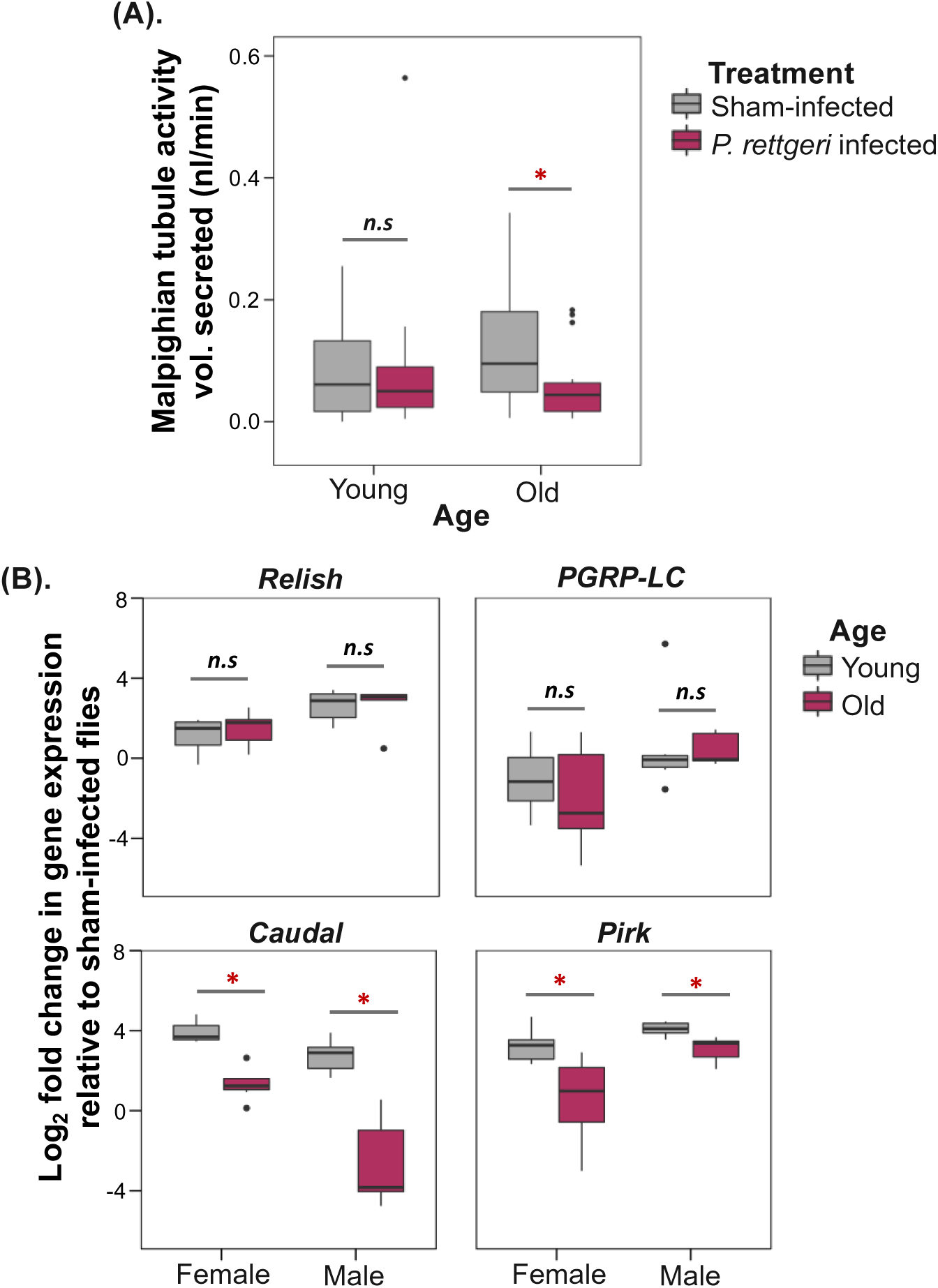
Ageing-associated immune dysregulation and immunopathology. **(A)** Malpighian tubule (MT) activity (n = 12-20 females/infection treatment/age-group), as a proxy for immunopathological damage, measured at 4-hours after infection with 0.1 OD of *P. rettgeri.* Statistically significant difference between groups are indicated by asterisk (*). **(B)** Expression of positive (*Relish, PGRP-LC*) and negative (*Caudal, Pirk*) regulators of Imd-pathway across sexes and age-groups after *P. rettgeri* infection, relative to an internal control *rp49* (n= Total 15-21 flies homogenized in Trizol in a group of 3/Infection treatment/ age-group/ sex-combination). ns = not significant

Finally, we also found ageing-associated downregulation of the major negative regulators of Imd-signalling such as *Caudal* & *Pirk* in older flies (Fig. 5B; Table S12), which has been previously linked to the production of toxic levels of AMP production, causing reduced lifespan, locomotor defects and extensive neurodegeneration (Kounatidis et al., 2017; Prakash et al., 2021). Based on these results, we speculate that the observed expansion of AMP repertoire with age is therefore most likely to represent suboptimal body condition, characterized by poorly regulated immune system and increased physiological costs.

## Discussion

Recent studies performed functional validation of *Drosophila* AMPs, revealing remarkable specificity and non-redundant interactions with subsets of pathogens that they target (Hanson et al., 2019). In the present work, we analysed these specific AMPs responses primarily as a function of ageing that alters the regulation and relative investment in immune responses (Khan et al., 2016, 2017). We used two bacterial entomopathogen *P. rettgeri* and *P. entomophila* to induce various level of AMP responses inside a fly host, ranging from a single AMP to pathway-specific expression [e.g., Imd vs Toll; (Hanson al., 2019)]. Further, although sex profoundly impacts the relative use of AMPs (Duneau et al., 2017), previous studies addressing AMP specificity have almost entirely focussed on males (Unckless et al., 2016; Hanson et al., 2019). We thus also included both males and females in our experiments to test the sex-specific effect of ageing on AMP functions. In fact, we showed that the efficiency of these AMP responses is strictly age-driven with high degree of sexual dimorphism. For example, the classic *Dpt*-driven protection against *P. rettgeri*, as shown by previous studies (Hanson et al., 2019), is only limited to young males, whereas females also needed other Imd-regulated AMPs. Although the reason is unclear, we speculate that multiple AMPs were needed possibly to compensate the inherently lower expression level of *Dpt* transcript in females than males (Duneau et al., 2017; also shown by Prakash, 2022). However, regardless of sex and pathogen, ageing led to a more drastic expansion of AMP repertoire— instead of deploying only canonical expression of Imd-responsive AMPs to counter Gram-negative bacterial infections, older males and females also used AMPs from Toll pathways.

Surprisingly, despite using more diverse AMPs, late-life expansion either did not confer any survival benefits (during *P. rettgeri* infection in older males) or was associated with survival costs (after *P. entomophila* infection). We thus speculate that the nonspecific use of AMPs with ageing was unnecessary, perhaps indicating an immune system failing to control over-reactive immune responses with potentially immunopathological effects (Stout-Delgado et al., 2009; Goldstein, 2010; Khan et al., 2017; Badinloo et al., 2018). This notion was further supported by reduced expression levels of negative regulators of immune responses such as *caudal* and *pirk* in older flies, which have been previously implicated in over-activating Imd-signalling and AMP expression. In addition, reduced renal function or Malpighian tubule activity in infected older flies suggested that expanded AMP repertoire might not be able to prevent the plausible physiological costs of bacterial infection. Although not verified experimentally, we suspect a causal role of overactivated AMPs here. This is because (a) overactive and simultaneously expressed multiple AMPs can impose cytotoxic effects (Badinloo et al., 2018), and (b) reduced Malpighian tubule activity is already a known manifestation of immunopathological costs caused by overactive insect immune components (Sadd and Siva-Jothy, 2006; Khan et al., 2017), reducing fitness by accumulating toxic metabolites (Li et al., 2020).

We also note an alternative possibility where age-specific increase in AMP expression could have been beneficial. For example, since ageing can lead to accumulation of diverse microbes in the body cavity (Ren et al., 2007; Arias-Rojas and Iatsenko, 2022), this might warrant the overexpression of multiple AMPs to tackle the antigenic diversity of many microbial species to maintain the health (Ren et al., 2007; Badinloo et al., 2018). Indeed, previous experiments have found that highly expressed Imd-responsive AMPs such as *CecA1* and *Dro* were needed to maintain health while extending the lifespan in *Drosophila* (Loch et al., 2017). However, benefits of non-specific, highly expressed immune responses may still not be able to outweigh the net costs of overreactive immune responses. In fact, detrimental effects of overreactive immunity with ageing has been supported by recent analyses linking weaker strength of purifying selection in older individuals and high frequency of non-synonymous and disease-causing mutations (Cheng and Kirkpatrick, 2021). This in turn can lead to poorly-regulated gene expression network in older animals with increased cancer risk in a range of species, including humans. Taken together, non-specific AMP responses with ageing is thus a more likely feature of a deregulated immune system of older individuals (Kounatidis et al., 2017).

Finally, the use of diverse array of AMP deletion mutants allowed us to capture enormous functional diversity of AMPs, revealing dynamic age- and sex-specific changes in their pathogen clearance ability. Older individuals showed increased divergence between individual AMPs vs their combined action (e.g., *Dpt* vs group-B mutants in older females), possibly indicating greater complexity associated with higher number of AMPs in use vs their various interactions. Although we did not find much evidence of synergism or additive effects between individual AMPs (but see the older males infected with *P. entomophila*), indispensability of each AMPs to maintain the fitness post-infection in older flies suggested the mutually non-exclusive and intertwined nature of their activity with ageing. We hope that these results will motivate future studies to investigate the deeper mechanistic details of nonspecific AMP function with ageing. Also, with growing importance of AMPs in developing novel antibiotics and autoimmune disease research, identifying age or sex as major sources of variability in AMP functions and fitness impacts might have significant importance for therapeutic and gerontological research.

## Acknowledgements

We thank Basabi Bagchi, Joy Bose, and Srijan Seal for feedback on the manuscript. We are grateful to Bruno Lemaitre and Mark A. Hanson for generously providing us the fly lines. We thank Srijan Seal, Katy M. Monteith, Raghav Pavan Thunga and Devshuvam Banerji for laboratory assistance.

## Author contribution

IK conceived the experiments; IK, AP, BS designed the experiments; AP, BS, and SS performed the experiments; AP, BS and IK analysed the data; IK and PV acquired the funding and provided resources and consumables. IK and AP drafted the manuscript with additional input and comments from BS, SS and PV. All authors agreed on the final version of the manuscript.

## SUPPLEMENTARY METHODS

### i. Bacterial culture preparations & systemic infections

We used two gram-negative bacterial pathogens *Providencia rettgeri* and *Pseudomonas entomophila*, with a broad host range infecting insects, nematodes, plants and also higher vertebrates (Siva-Jothy et al., 2018; Troha and Buchon, 2019). In fruit flies, *P. rettgeri* and *P. entomophila* infection can result in severe pathology, eventually causing death (Vodovar et al., 2005; Dieppois et al., 2015; Galac and Lazzaro, 2011). In brief, we used overnight grown cultures of *P. rettgeri* & *P. entomophila* at 37°C and 30°C respectively with shaking at 120 rpm obtained from pure bacterial isolates.

We used LB [Luria Bertani broth; see (Siva-Jothy et al., 2018) for recipe] as a growth media for the bacterial cultures. At the mid-log phase (OD_600_ = 0.95), we harvested the bacterial cells by centrifugation at 5000 rpm for 5 min at 4°C and re-suspended the bacterial pellet in 1X PBS (phosphate buffer saline). We adjusted the final inoculum to OD_600_ = 0.1 for *P. rettgeri* & 0.05 OD for *P. entomophila* for all systemic infections. Briefly, we pricked the adult flies (young 3-5 day old, old 25-28 day old) in the thorax region (Khalil et al., 2015) using 0.1 mm minutein needle (Fine science tools ltd.). The OD 0.1 *P. rettgeri* and 0.05 OD of *P. entomophila*. For mock infections, we replaced the bacterial cultures with sterile 1x PBS.

### ii. Bacterial load measurements

To test whether ageing reduces host’s ability to supress the bacterial growth, we quantified bacterial load in each of the fly lines at 24-hours (or 20-hours) after *Providencia rettgeri* (or *Pseudomonas entomophila*) infection across different age groups (3- vs 25-day-old) and sexes (as described in Prakash et al., 2021). First, we thoroughly washed individual flies (pooled in a group of 6 in 1.5ml centrifuge tube) with 70% ethanol for 30-seconds to remove the surface bacteria, followed by rinsing them twice with sterile distilled water. We then homogenized the surface sterilized individuals using a clean motorized pestle for approximately a minute in 180µl 1X PBS broth. We performed serial dilution of the fly homogenate up to 10^-5^-fold and added 4μL of the aliquot onto LB agar plate. We incubated the plates overnight for 18-hours at 30◦C and counted the resultant bacterial colonies. None of sham-infected control fly homogenates produced any colonies on LB agar plates.

### iii. Assay for the Malpighian tubule activity

Insect Malpighian tubules are functionally analogous to mammalian kidney, playing vital roles in mediating osmoregulation, detoxification, and excretion (Chapman, 1998; Li et al., 2020). Their activity is of particular importance during infection because of their increased vulnerability to damage during immune activation against target pathogens (Sadd and Siva-Jothy, 2006). In fact, previous studies have provided an *in vitro* functional estimate of the ability of isolated tubules to transport saline across the active cell wall into the tubule lumen to demonstrate a large reduction in MT function due to immunopathological damage associated with immune activation in mealworm beetles *Tenebrio molitor* (Sadd and Siva-Jothy, 2006; Khan et al., 2017). Here, we used Ramsay assay (as described in Dow et al., 1994; Li et al., 2020) to estimate the fluid transporting capacity of MTs harvested from experimental flies after 4-hours of 0.1 OD *P. rettgeri*, as a proxy for immunopathology owing to immune response. Briefly, the flies were dissected in Schneider’s insect medium where intact MTs were first detached from the gut. One end of tubules were immersed in a mixture of Schneider’s medium and insect saline (bathing buffer), whereas the other end was pulled out of the buffer and wrapped around a minutein pin (0.1 mm). We allowed the fluid to secrete for 3-hours from the common ureter, following which the volume of secreted fluid was quantified. We considered the volume of the secreted droplet to be negatively correlated with the degree of immunopathological harm to MTs during bacterial infection (Khan et al., 2017). We analysed the MT activity data using a generalised linear model best fitted to quasi-binomial distribution.

### iv. Gene expression studies

The expression of major Imd-pathway regulators was quantified by qRT-PCR for both young and older individuals, as described previously in Prakash et al. (2021). We randomly selected a subset of young and old iso-*w^1118^* flies infected with 0.1 OD *P. rettgeri* for RNA extraction at 24h post-infection. We randomly removed selected flies (3 flies per vial) at time point 24 hours post exposure to *P. rettgeri*. We then homogenised the whole flies into 1.5µl microcentrifuge tubes containing 80µl of TRIzol reagent (Ambion, Life Technologies) using sterile micro-pestles (15-21 replicate flies homogenized in a group of 3 flies/ sex/ age-groups). The tubes containing the homogenate were kept frozen at −80°C till we could extract the RNA following the manufacture’s protocol (Zymo research Ltd).

After confirming the purity of the eluted samples using a Nanodrop 2000 Spectrophotometer (version 3.8.1), we performed the reverse transcription (RT), where cDNA was synthesized from 2µl of the eluted RNA using M-MLV reverse transcriptase (Promega) and random hexamer primers, followed by 1: 1 dilution in nuclease free water. We then performed quantitative qRT-PCR using an Applied Biosystems machine with a Fast SYBR Green Master Mix (Invitrogen). For each replicate, we set up the 10µl reaction mix containing 1.5L of 1:1 diluted cDNA, 5µl of Fast SYBR Green Master Mix and a 3.5µl of a primer stock containing both forward and reverse primer at 1µM suspended in nuclease free water (final reaction concentration of each primer was 0.35µM; see Table S1 for qPCR primers used in this study). For each cDNA sample across gene of interests, we had two technical replicates. We calculated the mean threshold cycle (Ct) for the further analyses as described in Livak and Schmittgen (2001).

We analysed the gene expression data by first calculating the ΔCt value for the expression of gene of interest vs the house-keeping gene *rp49,* across sexes after *P. rettgeri* (or sham) infection. We then calculated the fold change in gene expression in infected flies relative to the sham-infected flies. We found the data to be non-normally distributed (confirmed using Shapiro-Wilks’s test). We thus log-transformed data to fit into a normal distribution and analysed the data using an ANOVA. For each gene of interest, we specified the model as: Fold-change~ Age, with age as fixed effects across genes and sexes separately.

## SUPPLEMENTARY FIGURES

**Figure S1.**
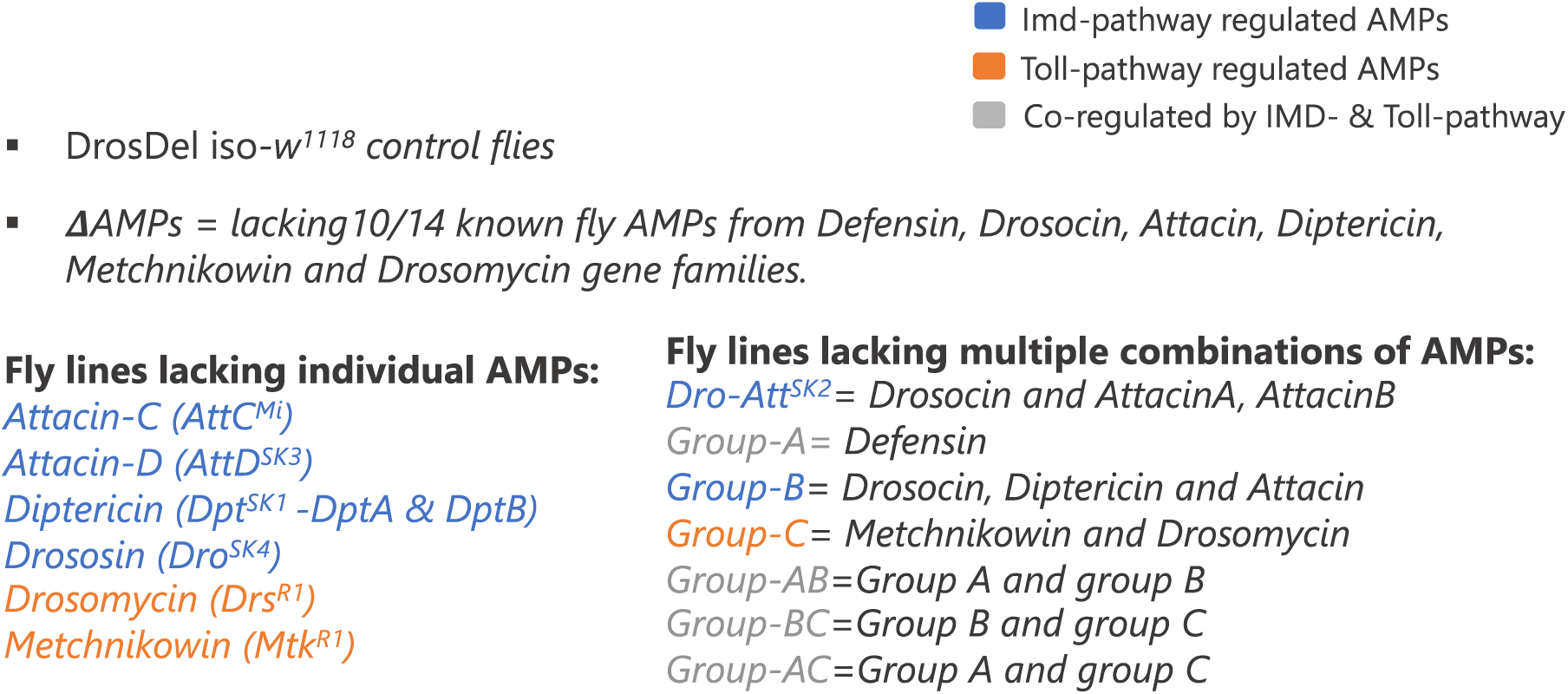
List of fly lines used in the study including background or control (iso-*w^1118^*), fly lines lacking single (individual) or multiple (combinations) of pathway-specific AMP knockouts that are Imd, Toll or co-regulated.

**Figure S2.**
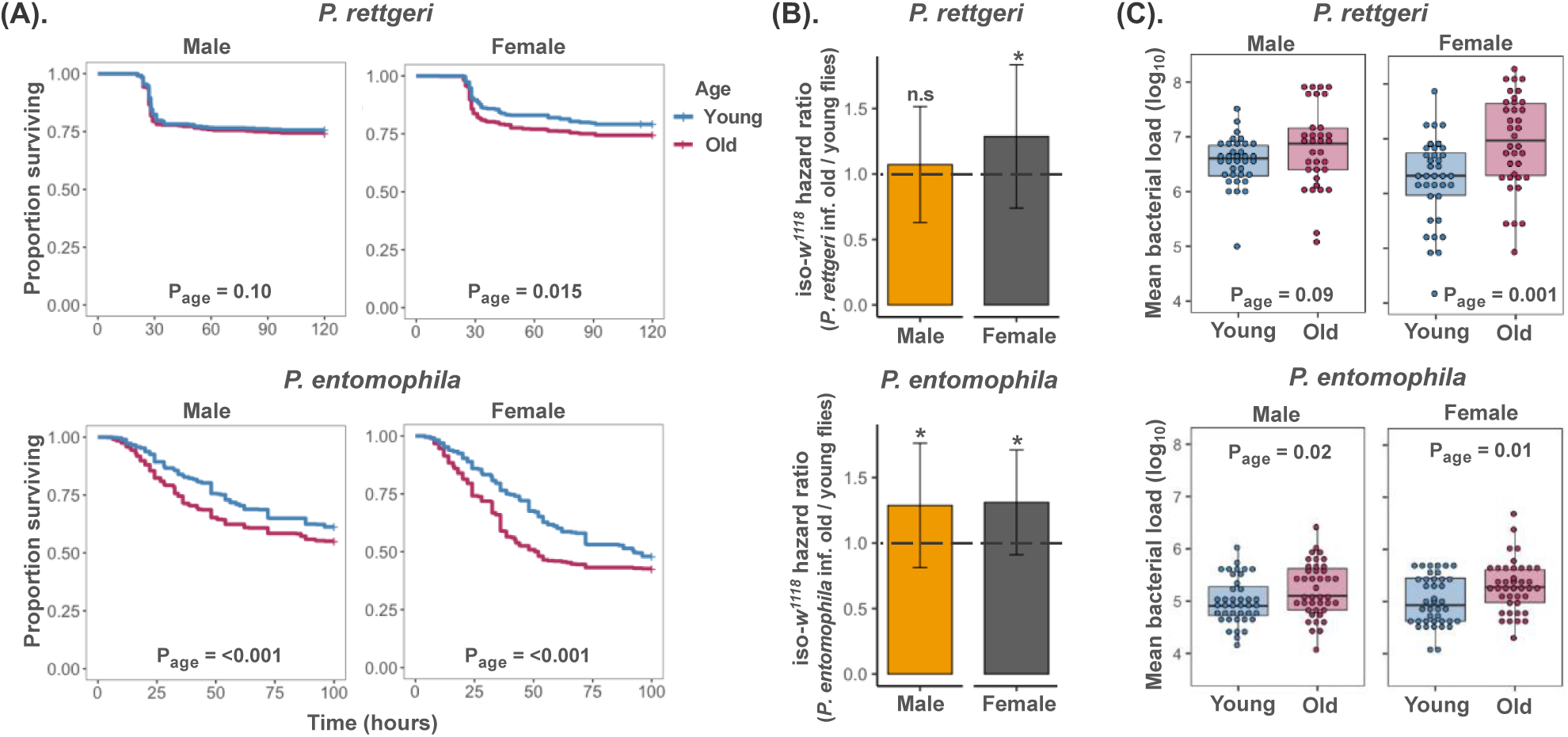
Post-infection survival and bacterial load data of iso-*w^1118^* flies across age-groups **(A)** Survival curves for iso-*w^1118^* male and females infected with *P. rettgeri* or *P. entomophila* across age-groups (n= 160-280 flies/infection treatment/ age-group/ sex/ replicate experiment). Survival data were analysed with a mixed effects Cox model [model: survival ~ age + (1|food vials + 1| replicate experiments), with ‘age’ as fixed effects and, ‘food vials’ and ‘replicate experiments’ as random effects. **(B).** Estimated hazard ratios for old vs young iso-*w^1118^* males and females. A hazard ratio significantly greater than 1 indicates higher susceptibility of older iso-*w^1118^* flies to infection than their younger counterparts. **(C).** Bacterial load measured as colony forming units (CFUs) after 24- (or 20-) hours of *P. rettgeri* (or *P. entomophila*) infection (n=8-15 replicate groups/treatment/age-groups/sex/ pathogen/ replicate experiments). Bacterial load data (log transformed) for each sexes were analysed using a generalized linear mixed model (GLMM) with Gamma distribution, using ‘age’ and ‘replicate experiment’ as a ‘fixed’ and ‘random effect’ respectively.

**Figure S3.**
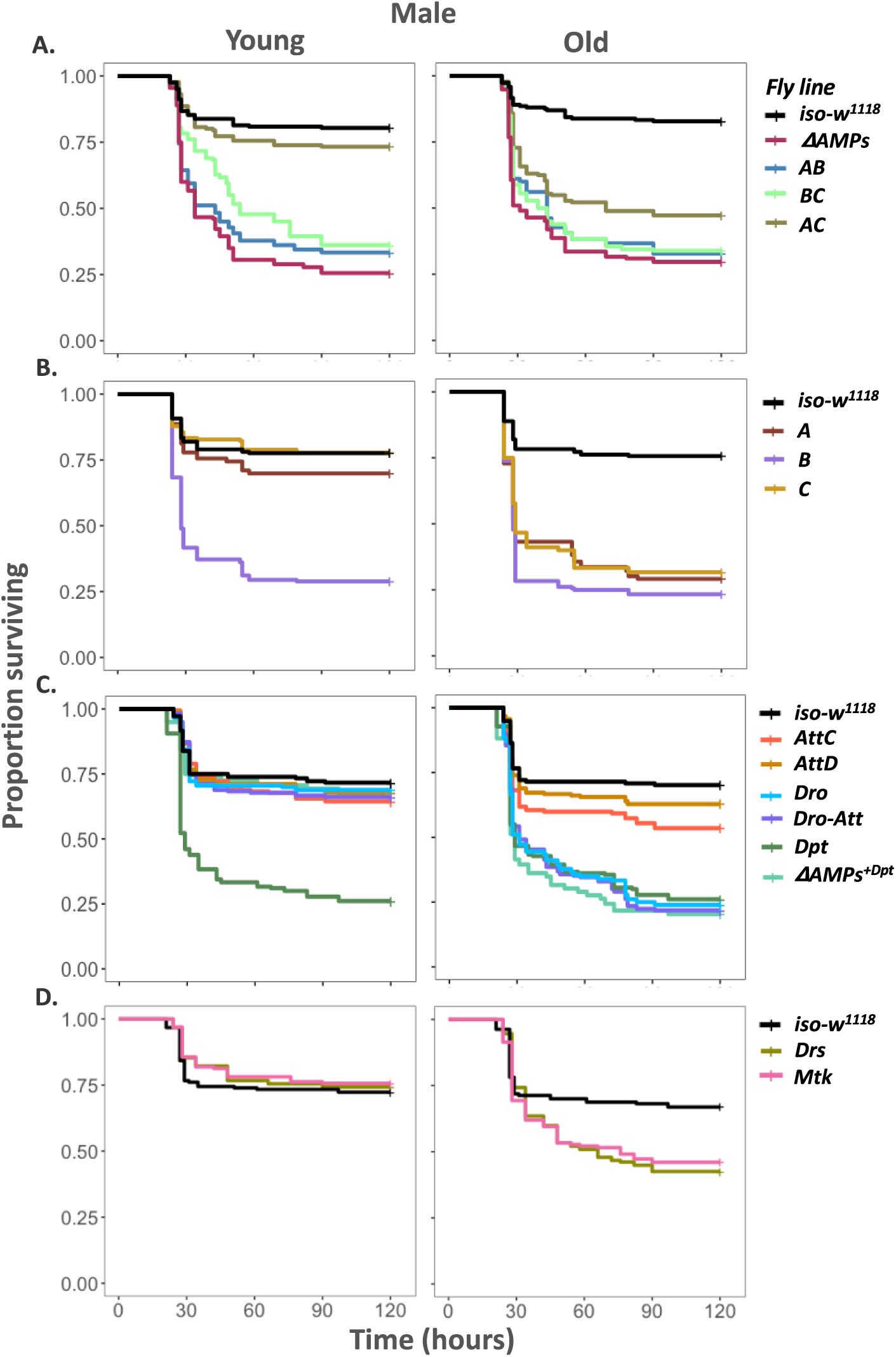
Survival curves for young and old males following *P. rettgeri* infection. Control iso-*w^1118^* flies vs **(A)** Combined mutants and *ΔAMPs*; **(B)** Pathway-specific mutants; **(C)** Single mutants lacking Imd-responsive AMPs (i.e., *Dpt, AttC, AttD, Dro*), Δ*AMPs^+Dpt^*and *Dro-Att* **(D)** Single mutants lacking Toll-responsive AMPs (e.g., *Drs & Mtk*) (n= 160-180 flies/infection treatment/ fly lines/age combination).

**Figure S4.**
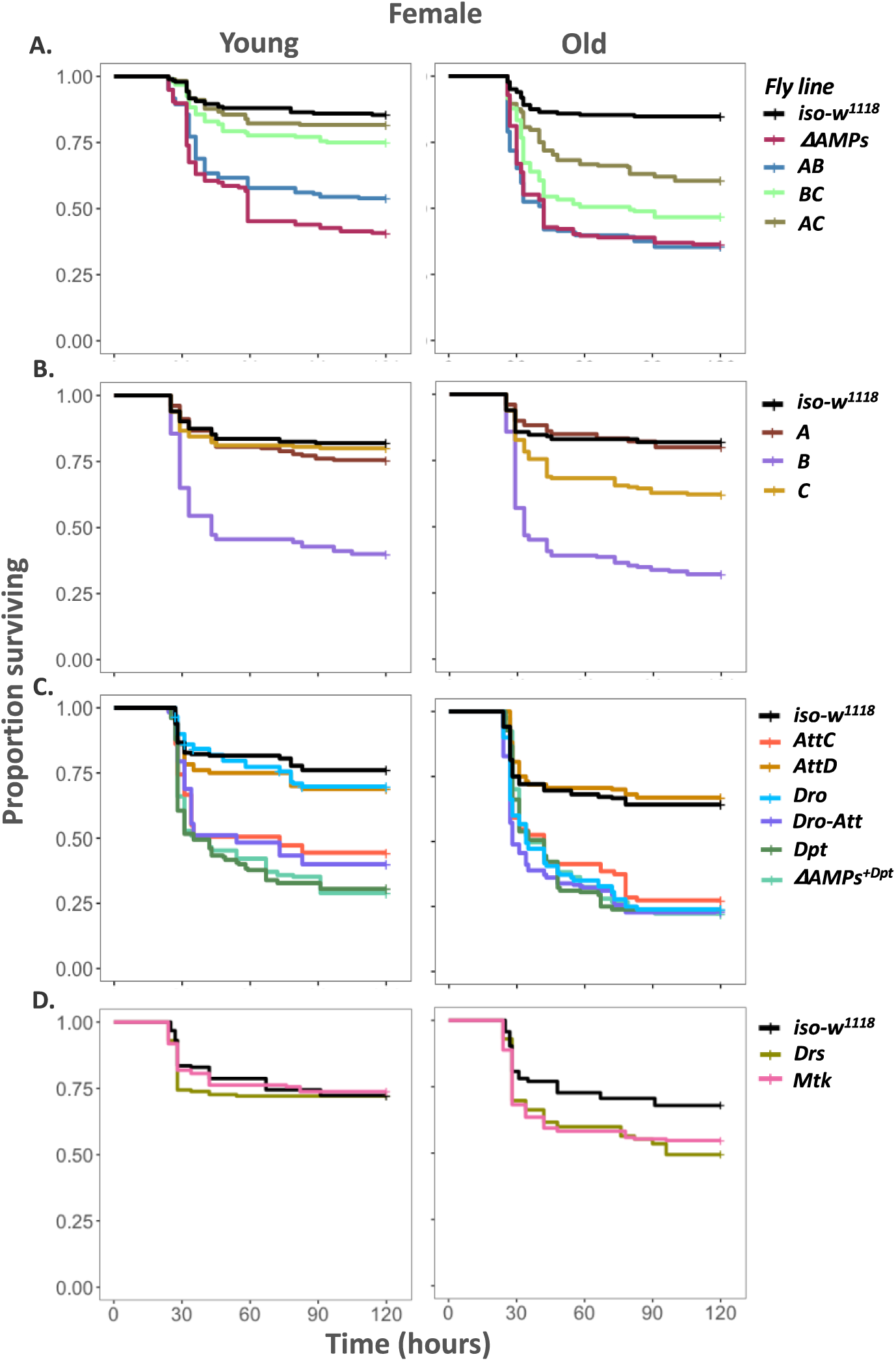
Survival curves for young and old females following *P. rettgeri* infection. Control iso-*w^1118^* flies vs **(A)** Combined mutants and *ΔAMPs*; **(B)** Pathway-specific mutants; **(C)** Single mutants lacking Imd-responsive AMPs (i.e., *Dpt, AttC, AttD, Dro*), Δ*AMPs^+Dpt^*and *Dro-Att* **(D)** Single mutants lacking Toll-responsive AMPs (e.g., *Drs & Mtk*). (n= 160-180 flies/infection treatment/ fly lines/age combination).

**Figure S5.**
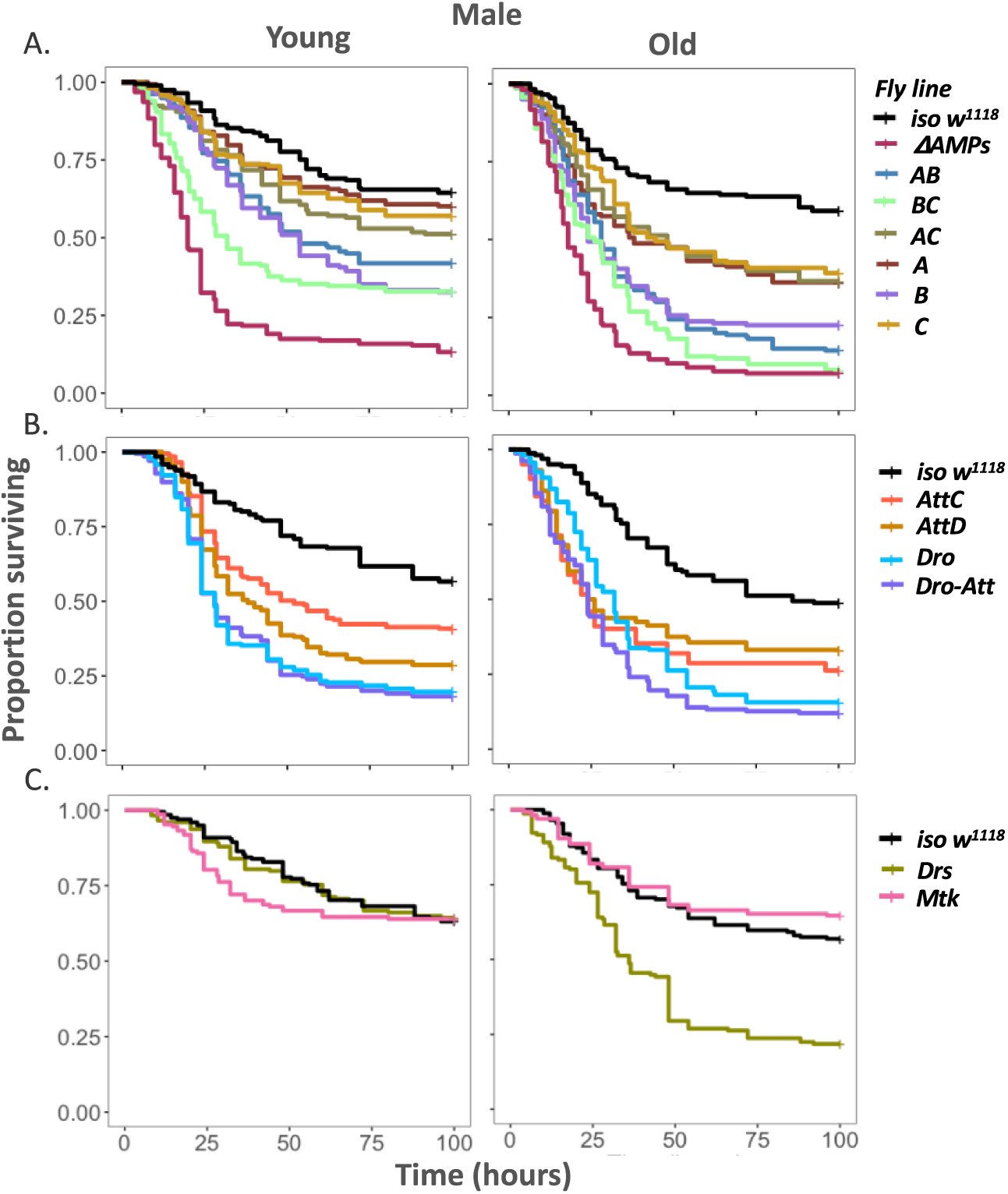
Survival curves for young and old males following *P. entomophila* infection. Control iso-*w^1118^* flies vs **(A)** Combined and pathway-specific mutants, and *ΔAMPs*; (B) Single mutants lacking Imd-responsive AMPs (i.e., *AttC, AttD, Dro), and Dro-Att;* **(C)** Single mutants lacking Toll-responsive AMPs (e.g., *Drs & Mtk*) (n= 180-280 flies/infection treatment/ fly lines/age combination).

**SI Figure 6.**
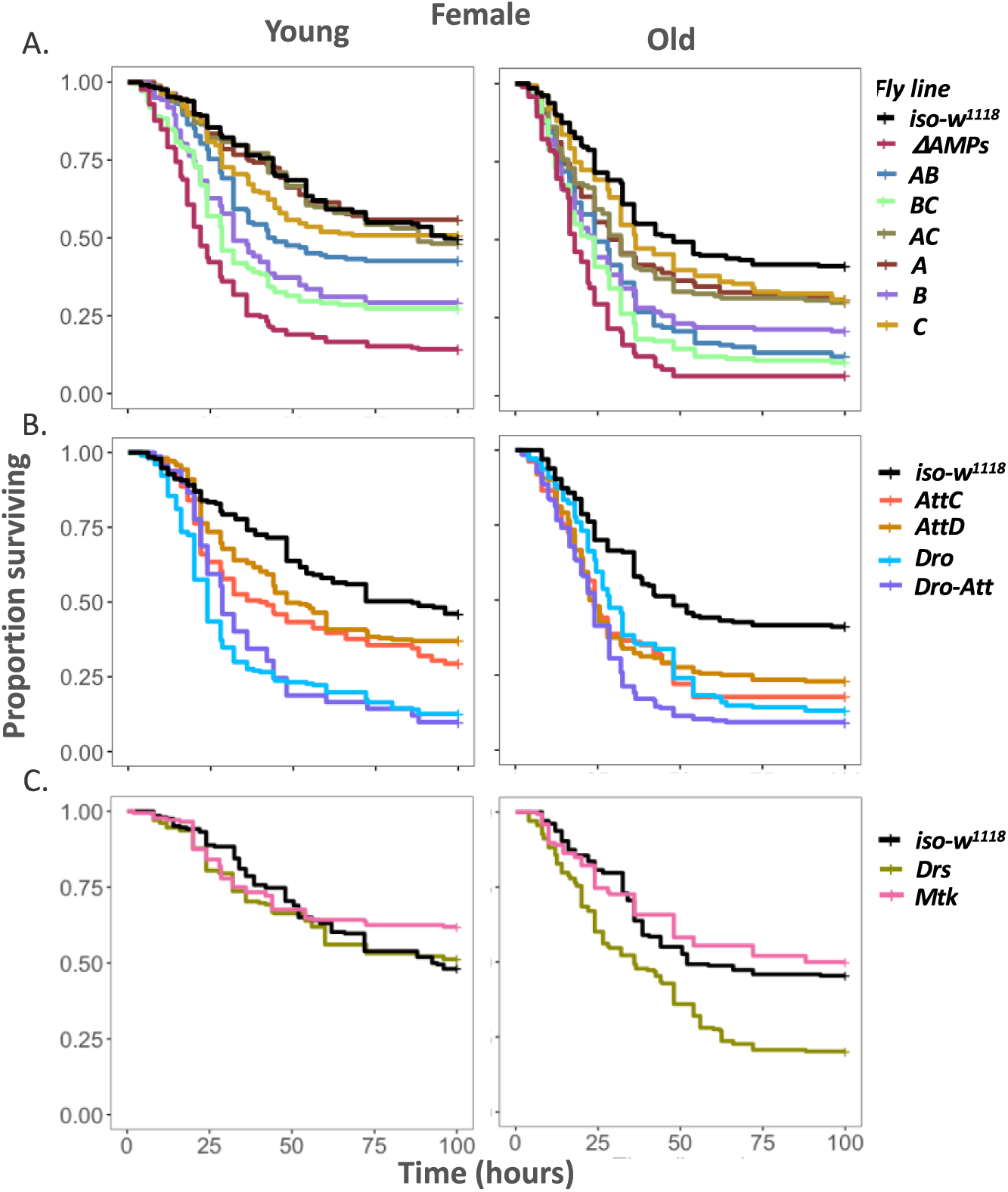
Survival curves for young and old females following *P. entomophila* infection. Control iso-*w^1118^* flies vs **(A)** Combined and pathway-specific mutants, and *ΔAMPs*; (B) Single mutants lacking Imd-responsive AMPs (i.e., *AttC, AttD, Dro), and Dro-Att;* **(C)** Single mutants lacking Toll-responsive AMPs (e.g., *Drs & Mtk*) (n= 180-280 flies/infection treatment/ fly lines/age combination).

## SUPPLEMENTARY TABLES

**Table S1:**
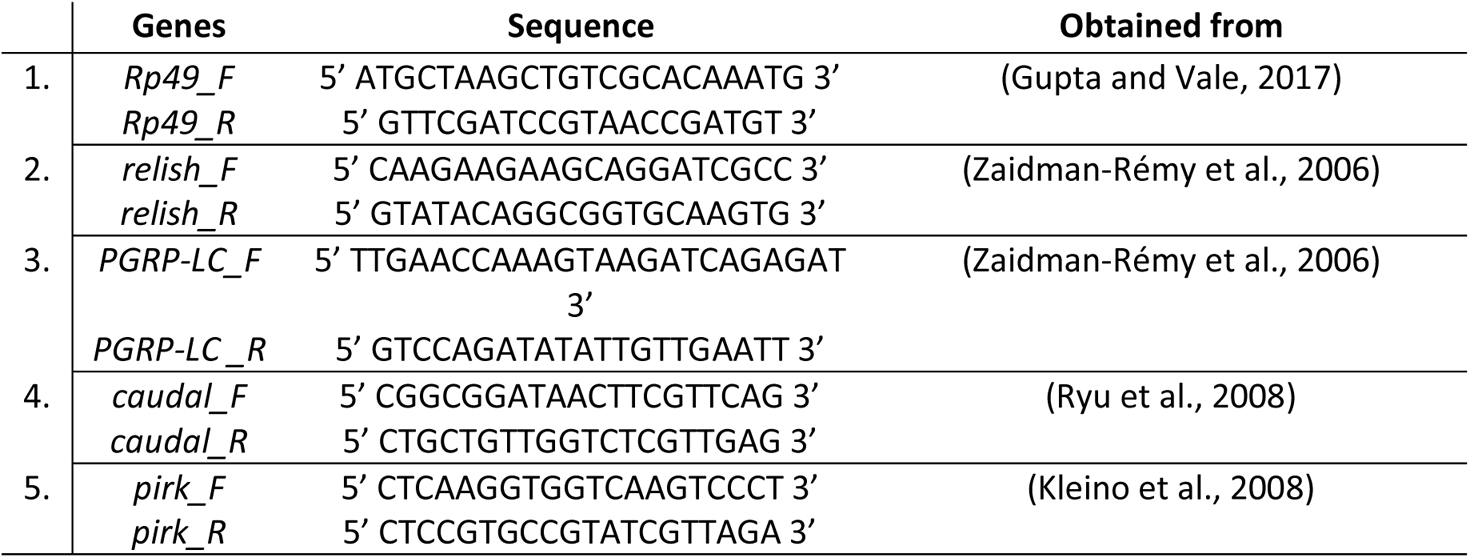
List of qPCR primers used in the study

**Table S2.**
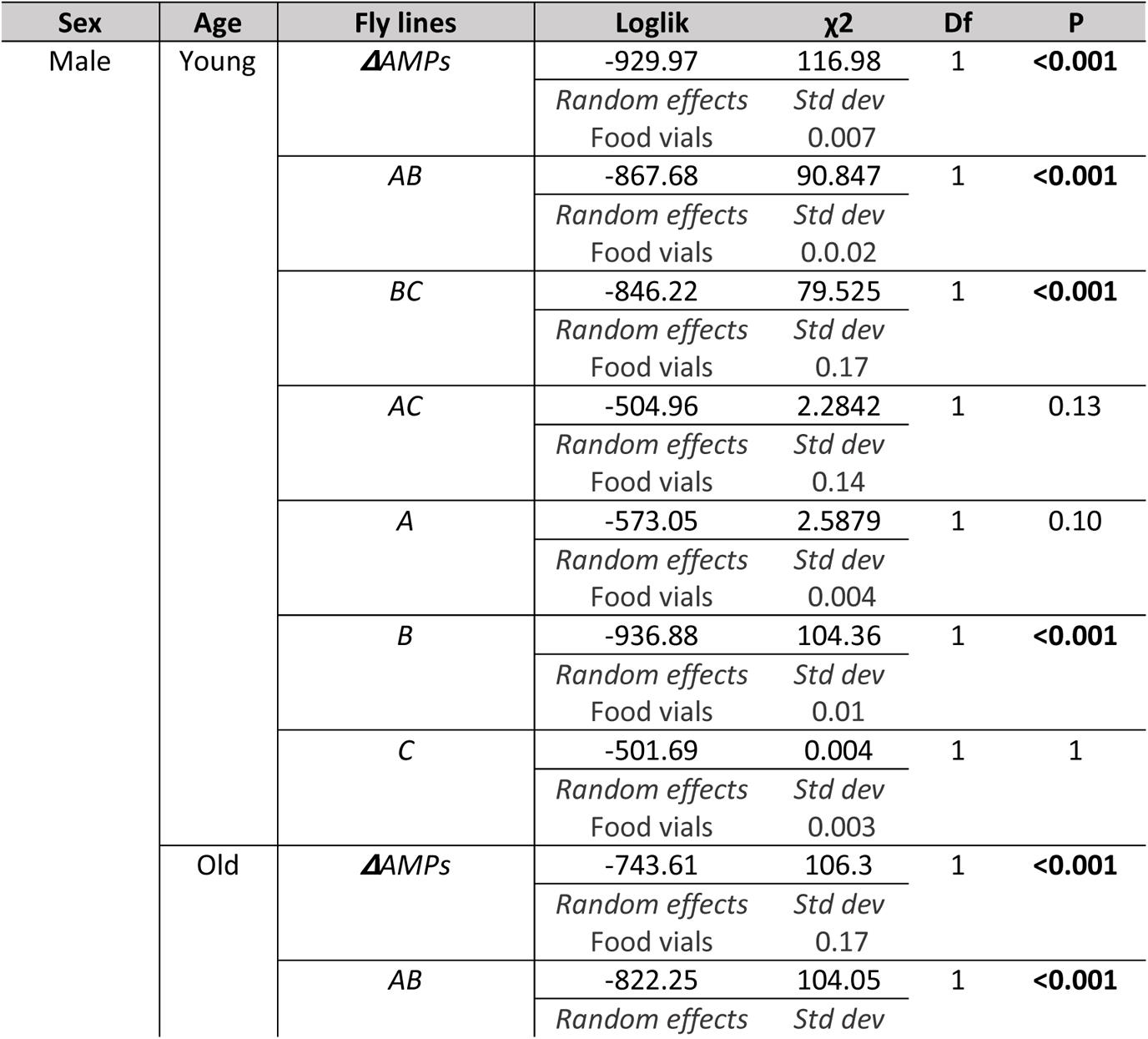
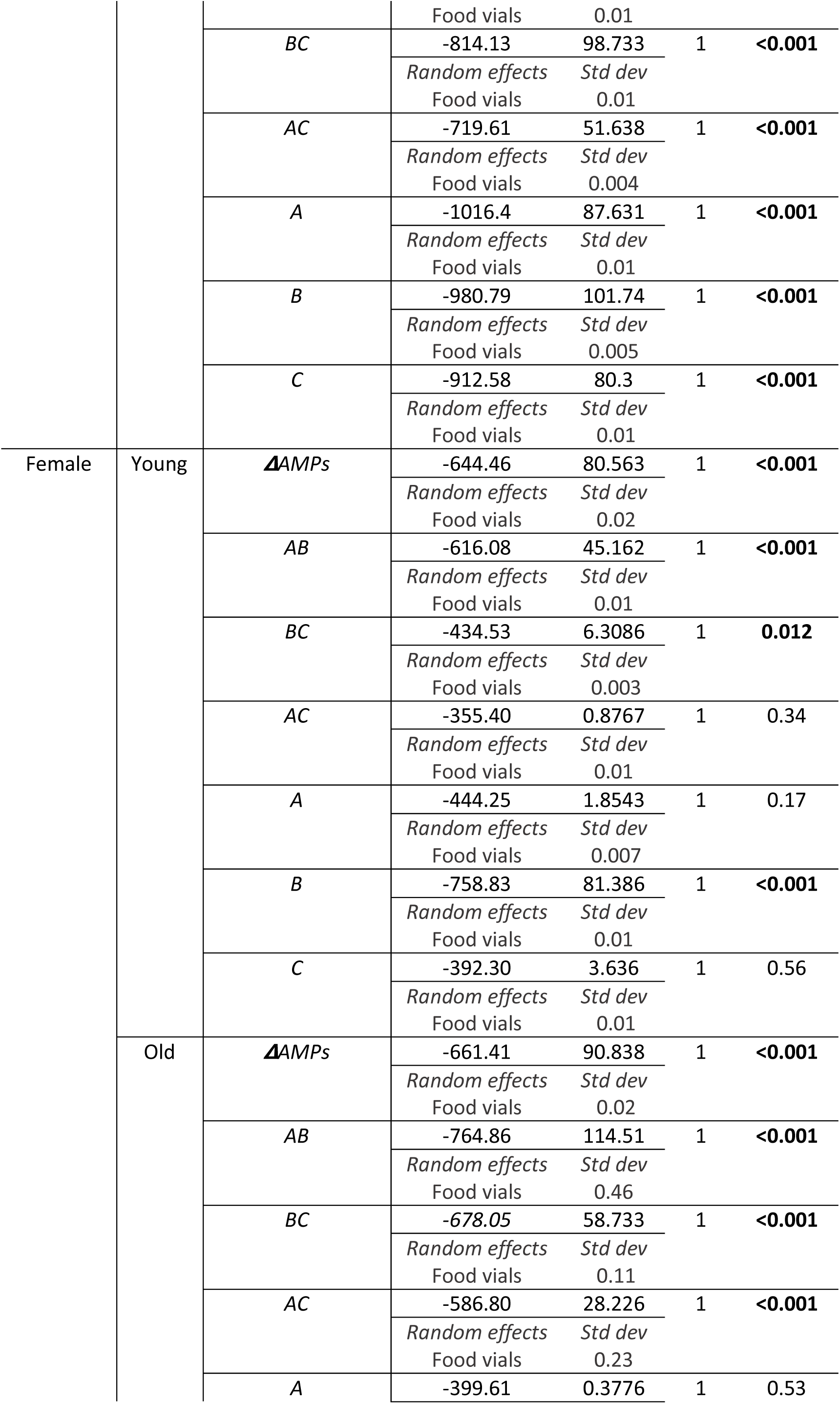
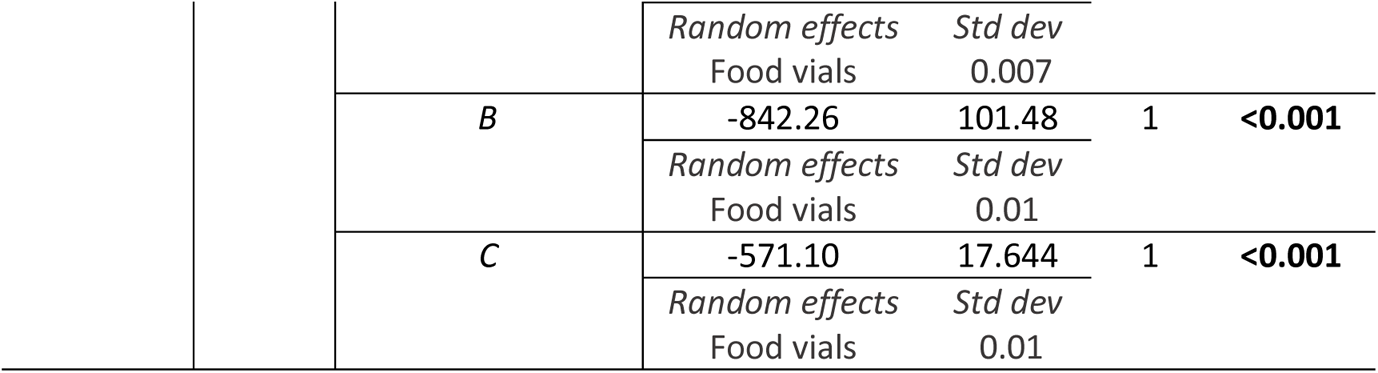
Summary of mixed effects Cox model to estimate the changes in the survival of individual fly lines with compound AMP mutations (i.e., A, B, C, AB, BC & AC), across sexes and age-groups, relative to control iso-*w^1118^* flies after *P. rettgeri* infection. We specified the model as: survival ~ fly line (individual mutant lines vs iso-*w^1118^*) + (1|food vials), with ‘fly line’ as a fixed effect and ‘replicate food vials’ as a random effect.

**Table S3.**
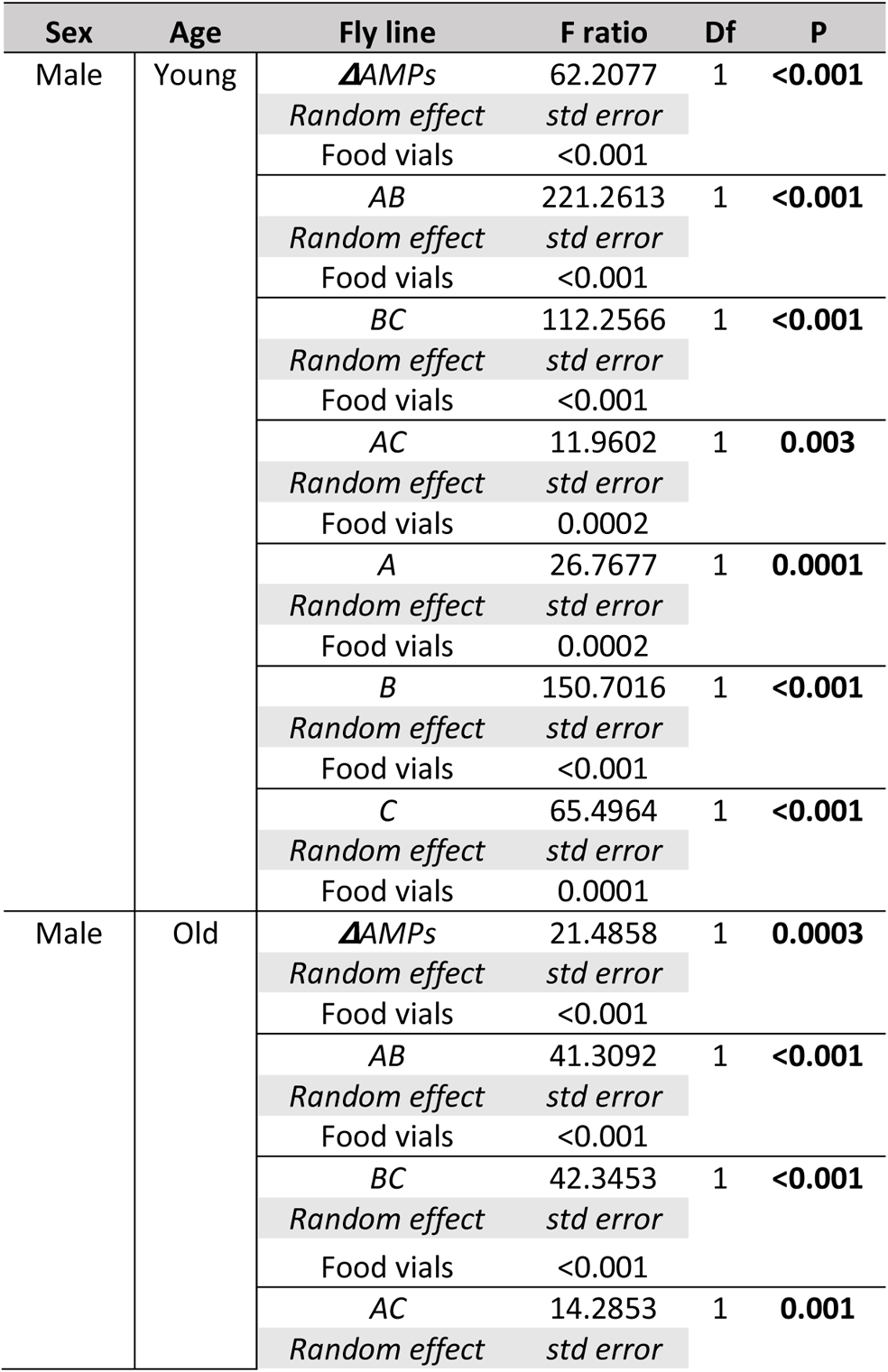
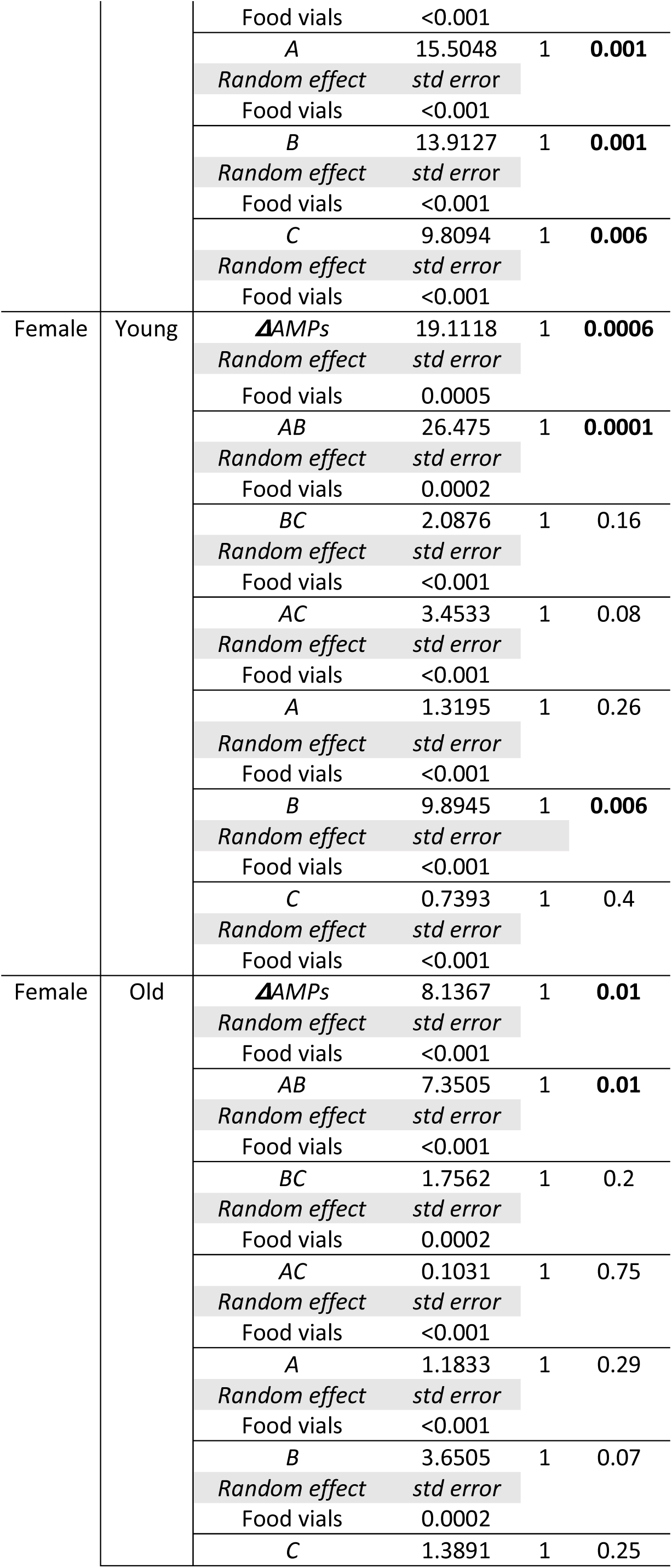
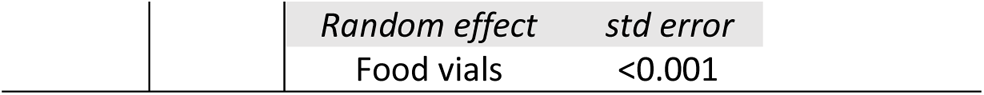
Summary of a generalized linear mixed model on log transformed bacterial load data (confirmed and fitted with a gamma distribution) for individual fly lines with compound mutations (i.e., A, B, C, AB, BC & AC) across sexes and age-groups, relative to control iso-*w^1118^* flies after *P. rettgeri* infection. We specified the model as: log transformed bacterial load ~ fly line (individual mutant lines vs iso-*w^1118^*), with ‘fly line’ as fixed and ‘replicate food vials’ as random effects across sexes and age-groups separately.

**Table S4.** **(A)** Summary of the mixed effects Cox model analyses on the age-specific survival data of iso-*w^1118^* males and females, infected with *P. rettgeri* and *P. entomophila*. For each pathogen species and sex, we specified the model as: Post-infection survival ~ Age + (1|Replicate experiment), with ‘age’ as a fixed effect and ‘replicate experiment’ as a random effect. **(B)** Summary of a generalized linear mixed-effects model, fitted to Gamma distribution, for age-specific bacterial load in iso-*w^1118^* males and females, after exposure to *P. rettgeri* and *P. entomophila* infection. For each pathogen species and sex, we specified the model as: Bacterial load ~ Age + (1|Food vial) + (1|Replicate experiment), with ‘age’ as a fixed effect, ‘food vials’ and ‘replicate experiment’ as random effects.

| A. Pathogen | Sex | Source | Loglik | Chi sq. | Df | P |
| --- | --- | --- | --- | --- | --- | --- |
| *P. rettgeri* | Male | Age | -2617.6 | 2.6135 | 1 | 0.10 |
|  |  | Random effect | Std dev |  |  |  |
|  |  | Food vials | 0.14 |  |  |  |
|  |  | Replicate experiment | 0.16 |  |  |  |
|  | Female | Age | -2505.6 | 5.8313 | 1 | **0.015** |
|  |  | Random effect | Std dev |  |  |  |
|  |  | Food vials | 0.07 |  |  |  |
|  |  | Replicate experiment | 0.17 |  |  |  |
| *P. entomophila* | Male | Age | -3083.8 | 24.269 | 1 | **<0.001** |
|  |  | Random effect | Std dev |  |  |  |
|  |  | Food vials | 0.27 |  |  |  |
|  |  | Replicate experiment | 0.20 |  |  |  |
|  | Female | Age | -4508.0 | 23.584 | 1 | **<0.001** |
|  |  | Random effect | Std dev |  |  |  |
|  |  | Food vials | 0.19 |  |  |  |
|  |  | Replicate experiment | 0.20 |  |  |  |

| B. Pathogen | Sex | Source | F ratio | Df | p |
| --- | --- | --- | --- | --- | --- |
| *P. rettgeri* | Male | Age | 2.956 | 1 | 0.09 |
|  |  | Random effect | Std dev. |  |  |
|  |  | Replicate experiment | 0.0009 |  |  |
|  | Female | Age | 12.56 | 1 | **0.0007** |
|  |  | Random effect | Std dev. |  |  |
|  |  | Replicate experiment | 0.0002 |  |  |
| *P. entomophila* | Male | Age | 5.147 | 1 | **0.026** |
|  |  | Random effect | Std dev. |  |  |
|  |  | Replicate experiment | 0.001 |  |  |
|  | Female | Age | 6.628 | 1 | **0.012** |
|  |  | Random effect | Std dev. |  |  |
|  |  | Replicate experiment | 0.0003 |  |  |

**Table S5.** Summary of the mixed effects Cox model analyses to estimate the changes in the survival of individual Imd- and Toll-pathway specific mutants, Δ*AMPs^+Dpt^*and *Dro-Att* across sexes and age-groups, relative to control iso-*w^1118^*flies after *P. rettgeri* infection. For each fly line across sexes and age-groups, we specified the model as: survival ~ fly line (individual fly lines vs iso-*w^1118^*) + (1|food vials), with ‘fly line’ as a fixed effect and ‘replicate food vials’ as a random effect.

| Sex | Age | Mutants | Loglik | $\chi^2$ | Df | P |
| --- | --- | --- | --- | --- | --- | --- |
| Male | Young | AttC | -655.69 | 1.377 | 1 | 0.24 |
|  |  |  | Random effects | Std dev |  |  |
|  |  |  | Food vials | 0.003 |  |  |
|  |  | AttD | -623.08 | 0.4223 | 1 | 0.51 |
|  |  |  | Random effects | Std dev |  |  |
|  |  |  | Food vials | 0.011 |  |  |
|  |  | Dro | -612.05 | 0.3547 | 1 | 0.55 |
|  |  |  | Random effects | Std dev |  |  |
|  |  |  | Food vials | 0.01 |  |  |
|  |  | Dpt | -984.71 | 81.284 | 1 | <0.001 |
|  |  |  | Random effects | Std dev |  |  |
|  |  |  | Food vials | 0.008 |  |  |
| | | ΔAMPS<sup>+Dpt</sup> | -582.66 | 0.8597 | 1 | 0.35 |
|  |  |  | Random effects | Std dev |  |  |
|  |  |  | Food vials | 0.003 |  |  |
|  |  | Dro-Att | -639.45 | 0.8147 | 1 | 0.36 |
|  |  |  | Random effects | Std dev |  |  |
|  |  |  | Food vials | 0.003 |  |  |
|  |  | Drs | -530.18 | 0.4169 | 1 | 0.51 |
|  |  |  | Random effects | Std dev |  |  |
|  |  |  | Food vials | 0.003 |  |  |
|  |  | Mtk | -512.44 | 0.827 | 1 | 0.36 |
|  |  |  | Random effects | Std dev |  |  |
|  |  |  | Food vials | 0.004 |  |  |
| Male | Old | AttC | -652.03 | 9.6893 | 1 | 0.001 |
|  |  |  | Random effects | Std dev |  |  |
|  |  |  | Food vials | 0.03 |  |  |
|  |  | AttD | -623.07 | 2.5085 | 1 | 0.11 |
|  |  |  | Random effects | Std dev |  |  |
|  |  |  | Food vials | 0.005 |  |  |
|  |  | Dro | -967.92 | 65.577 | 1 | <0.001 |
|  |  |  | Random effects | Std dev |  |  |
|  |  |  | Food vials | 0.004 |  |  |
|  |  | Dpt | -949.55 | 62.044 | 1 | <0.001 |
|  |  |  | Random effects | Std dev |  |  |
|  |  |  | Food vials | 0.004 |  |  |
| | | ΔAMPS<sup>+Dpt</sup> | -861.39 | 75.056 | 1 | <0.001 |
|  |  |  | Random effects | Std dev |  |  |
|  |  |  | Food vials | 0.006 |  |  |
|  |  | Dro-Att | -979.95 | 69.954 | 1 | <0.001 |
|  |  |  | Random effects | Std dev |  |  |
|  |  |  | Food vials | 0.004 |  |  |
|  |  | Drs | -815.79 | 12.942 | 1 | 0.0003 |
|  |  |  | Random effects | Std dev |  |  |
|  |  |  | Food vials | 0.003 |  |  |
|  |  | Mtk | -773.24 | 10.206 | 1 | 0.001 |
|  |  |  | *Random effects*<br>Food vials | *Std dev*<br>0.004 |  |  |
| Female | Young | *AttC* | -789.20 | 39.231 | 1 | **<0.001** |
|  |  |  | *Random effects*<br>Food vials | *Std dev*<br>0.006 |  |  |
|  |  | *AttD* | -566.75 | 2.133 | 1 | 0.14 |
|  |  |  | *Random effects*<br>Food vials | *Std dev*<br>0.003 |  |  |
|  |  | *Dro* | -548.26 | 3.3923 | 1 | 0.065 |
|  |  |  | *Random effects*<br>Food vials | *Std dev*<br>0.01 |  |  |
|  |  | *Dpt* | -903.67 | 76.398 | 1 | **<0.001** |
|  |  |  | *Random effects*<br>Food vials | *Std dev*<br>0.003 |  |  |
| | | ΔAMPs<sup>+Dpt</sup> | -829.03 | 73.907 | 1 | **<0.001** |
|  |  |  | *Random effects*<br>Food vials | *Std dev*<br>0.004 |  |  |
|  |  | *Dro-Att* | -827.71 | 48.025 | 1 | **<0.001** |
|  |  |  | *Random effects*<br>Food vials | *Std dev*<br>0.01 |  |  |
|  |  | *Drs* | -573.33 | 0.0986 | 1 | 0.75 |
|  |  |  | *Random effects*<br>Food vials | *Std dev*<br>0.004 |  |  |
|  |  | *Mtk* | -536.23 | 0.0158 | 1 | 0.89 |
|  |  |  | *Random effects*<br>Food vials | *Std dev*<br>0.004 |  |  |
| Female | Old | *AttC* | -1117.7 | 46.083 | 1 | **<0.001** |
|  |  |  | *Random effects*<br>Food vials | *Std dev*<br>0.005 |  |  |
|  |  | *AttD* | -717.91 | 0.9219 | 1 | 0.33 |
|  |  |  | *Random effects*<br>Food vials | *Std dev*<br>0.007 |  |  |
|  |  | *Dro* | -1150.0 | 55.462 | 1 | **<0.001** |
|  |  |  | *Random effects*<br>Food vials | *Std dev*<br>0.003 |  |  |
|  |  | *Dpt* | -1152.8 | 49.958 | 1 | **<0.001** |
|  |  |  | *Random effects*<br>Food vials | *Std dev*<br>0.003 |  |  |
| | | ΔAMPs<sup>+Dpt</sup> | -1059.5 | 46.363 | 1 | **<0.001** |
|  |  |  | *Random effects*<br>Food vials | *Std dev*<br>0.003 |  |  |
|  |  | *Dro-Att* | -1155.2 | 65.681 | 1 | **<0.001** |
|  |  |  | *Random effects*<br>Food vials | *Std dev*<br>0.003 |  |  |
|  |  | *Drs* | -824.05 | 11.458 | 1 | **0.0007** |
|  |  |  | *Random effects*<br>Food vials | *Std dev*<br>0.004 |  |  |
|  |  | *Mtk* | -771.29 | 7.4092 | 1 | **0.006** |
|  |  |  | *Random effects*<br>Food vials | *Std dev*<br>0.004 |  |  |

**Table S6.**
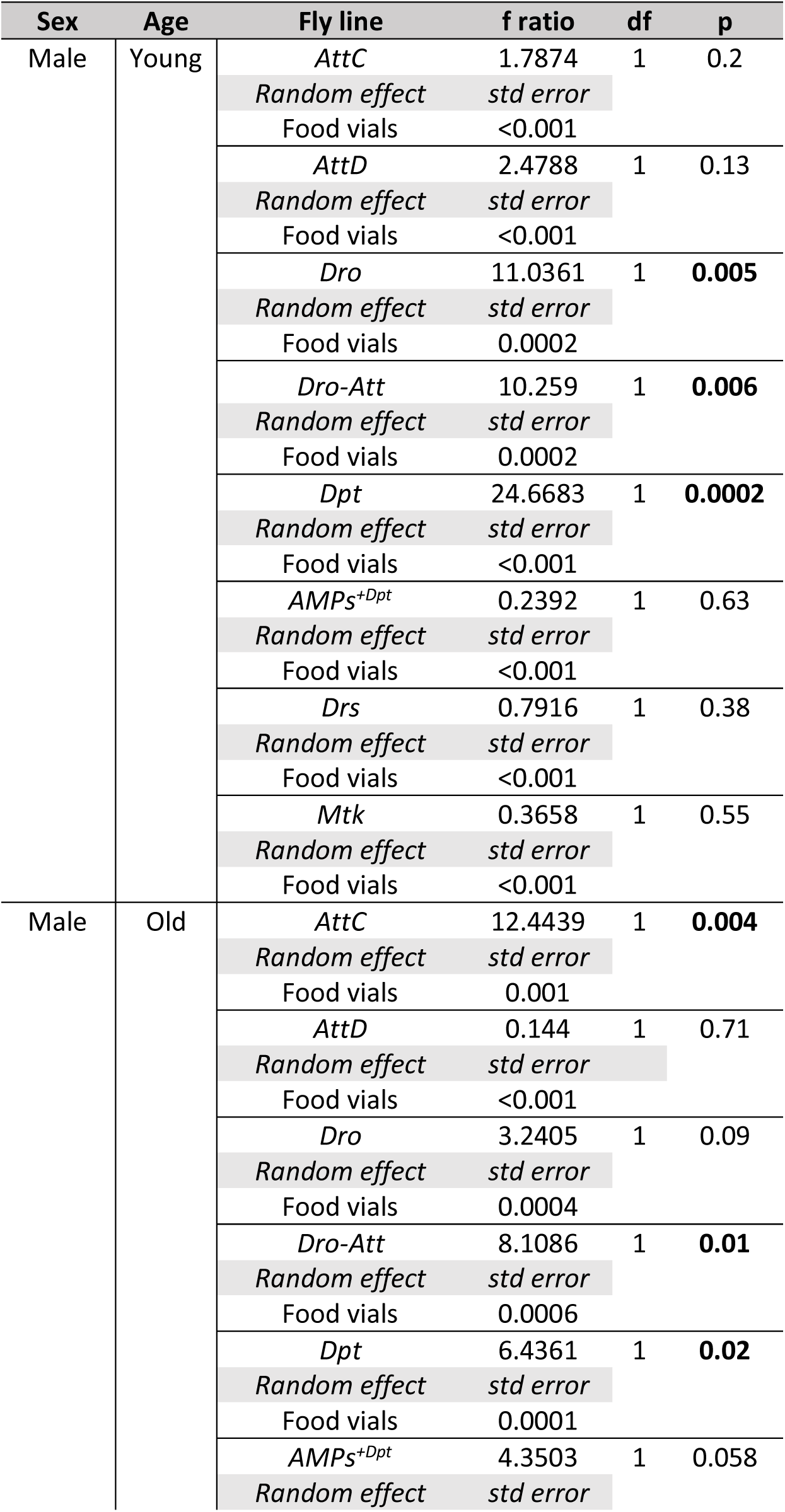
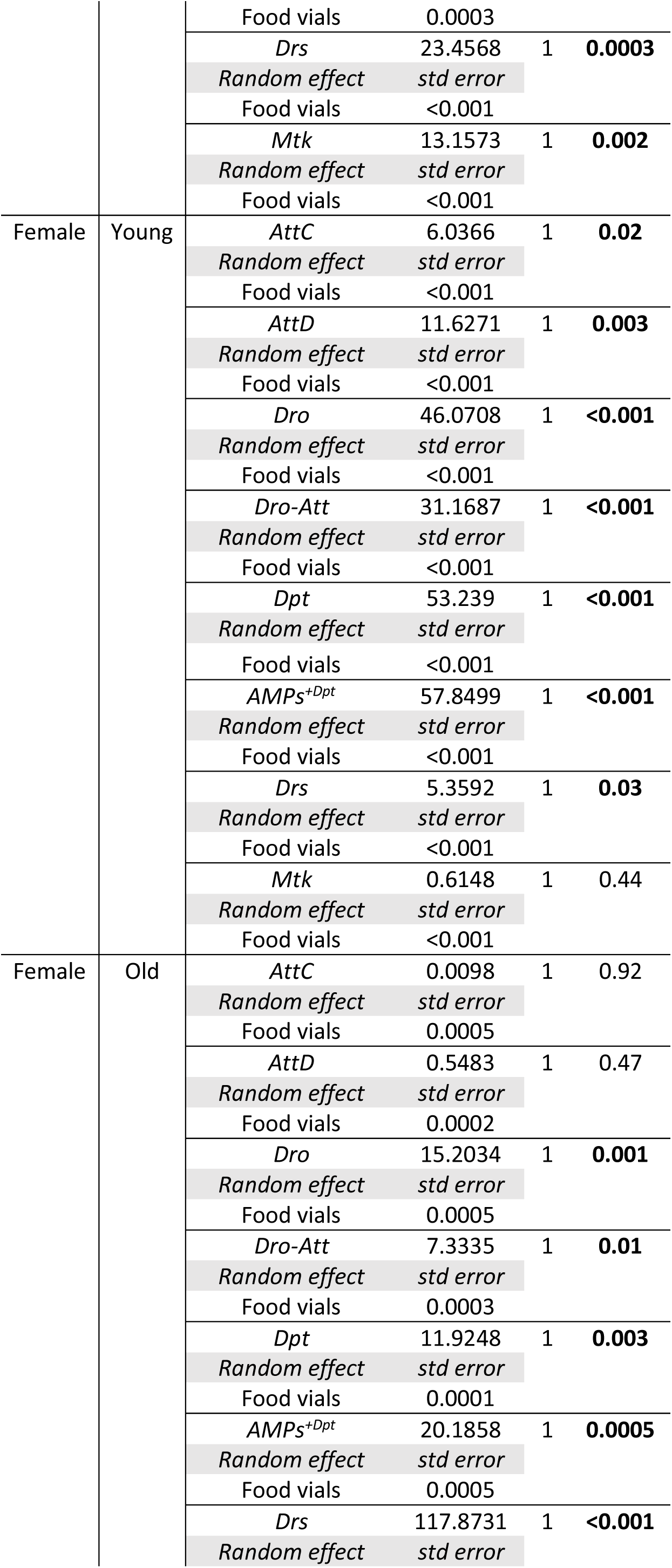
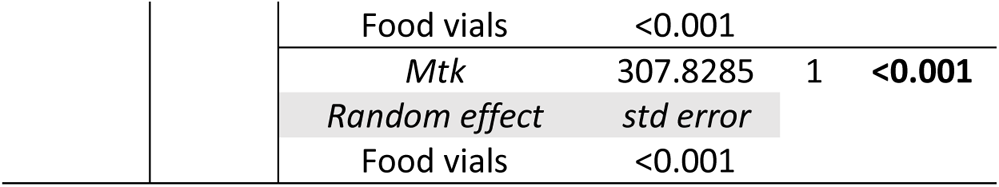
Summary of a generalized linear mixed model on log transformed bacterial load data (confirmed and fitted with a gamma distribution) for individual fly lines with Imd- & Toll-pathway specific, and *Dro-Att* mutations across sexes and age-groups, relative to control iso-*w^1118^* flies after *P. rettgeri* infection. We specified the model as: log transformed bacterial load ~ fly line (individual mutant lines vs iso-*w^1118^*) with ‘fly line’ and ‘replicate food vials’ as fixed and random effects across sexes and age-groups separately.

**Table S7.**
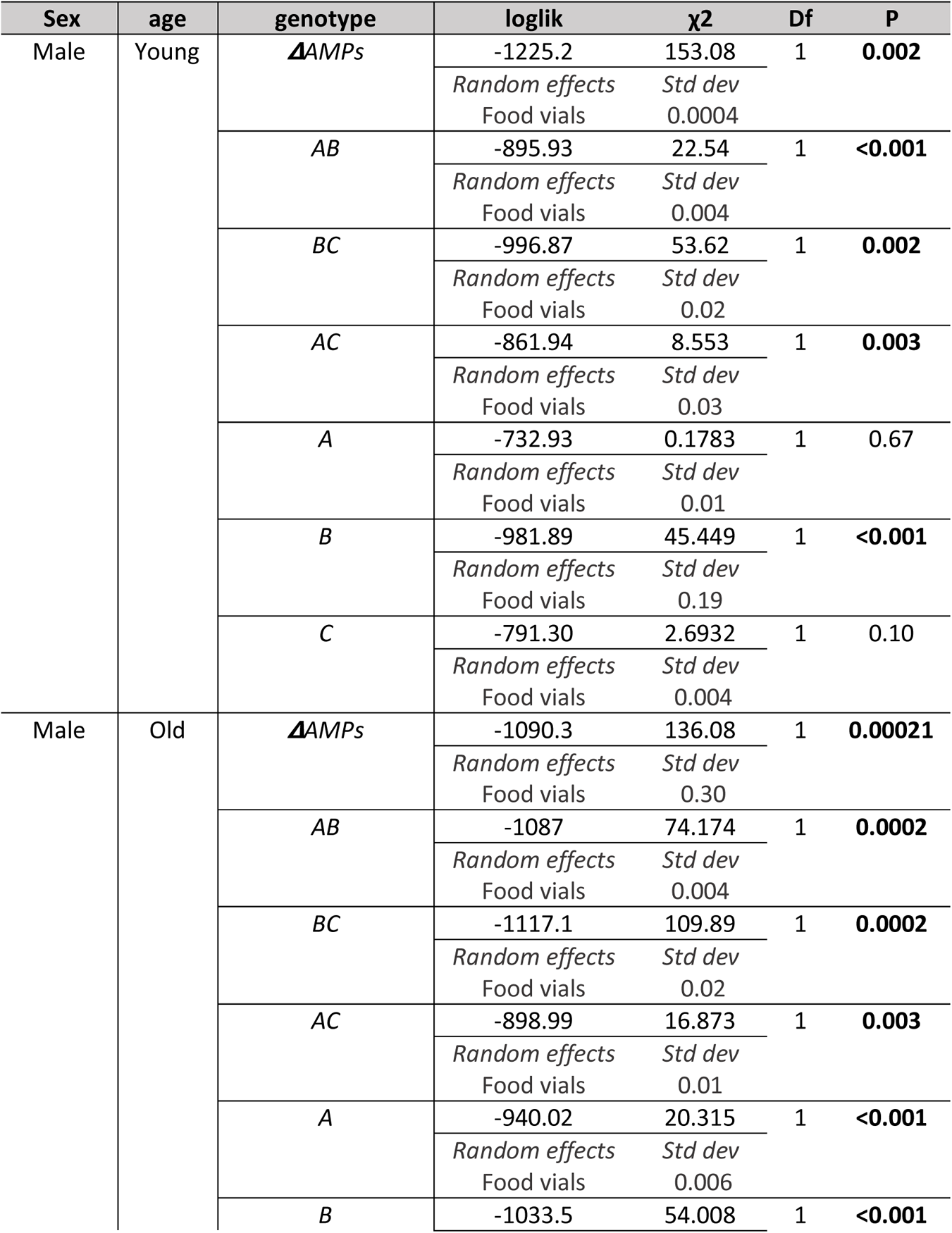
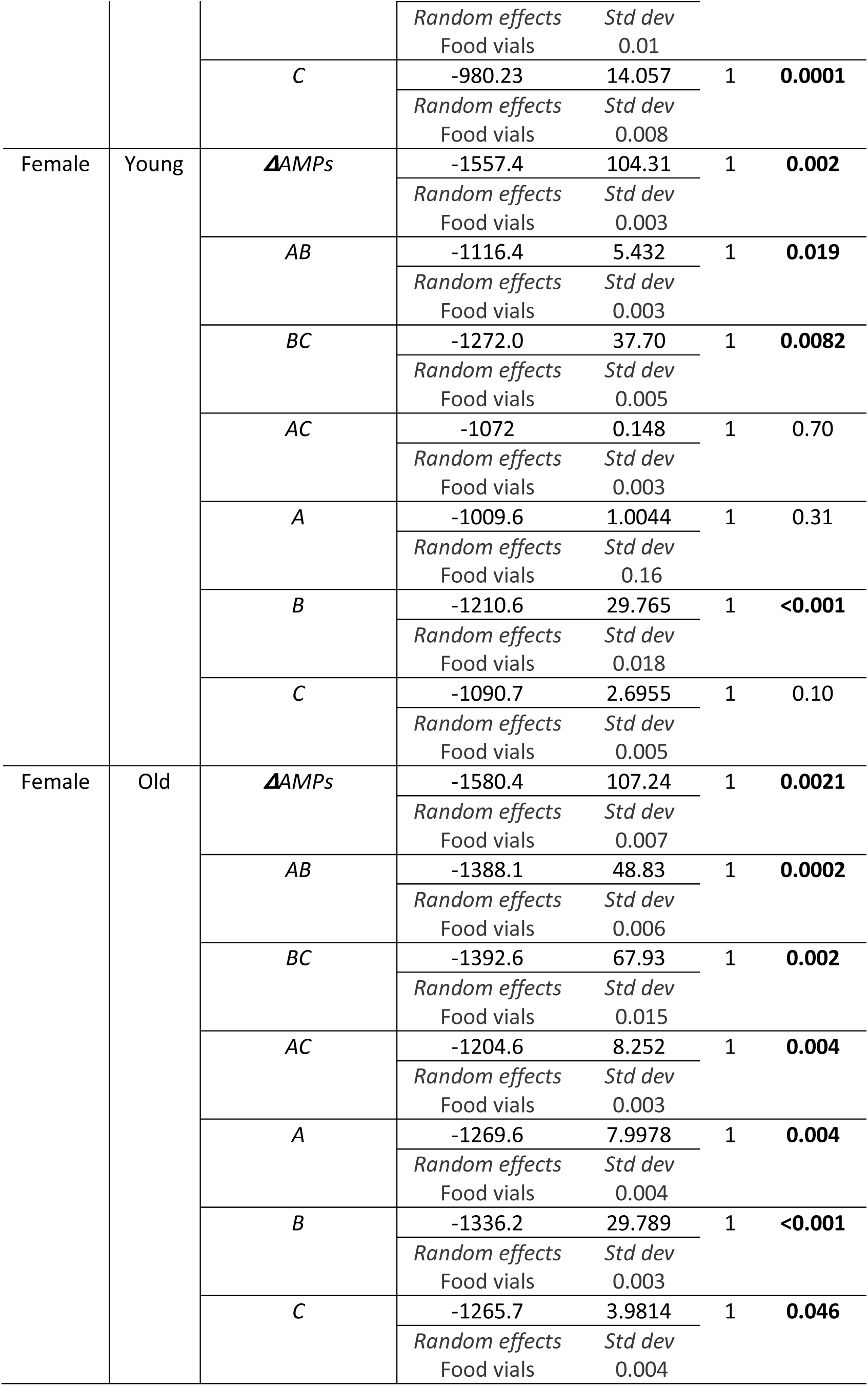
Summary of mixed effects Cox model to estimate the changes in the survival of individual fly lines with compound AMP mutations (i.e., A, B, C, AB, BC & AC), across sexes and age-groups, relative to control iso-*w^1118^* flies after *P. entomophila* infection. We specified the model as: survival ~ fly line (individual mutant lines vs iso-*w^1118^*) + (1|Food vials), with ‘fly line’ as a fixed effect and ‘replicate food vials’ as a random effect.

**Table S8.**
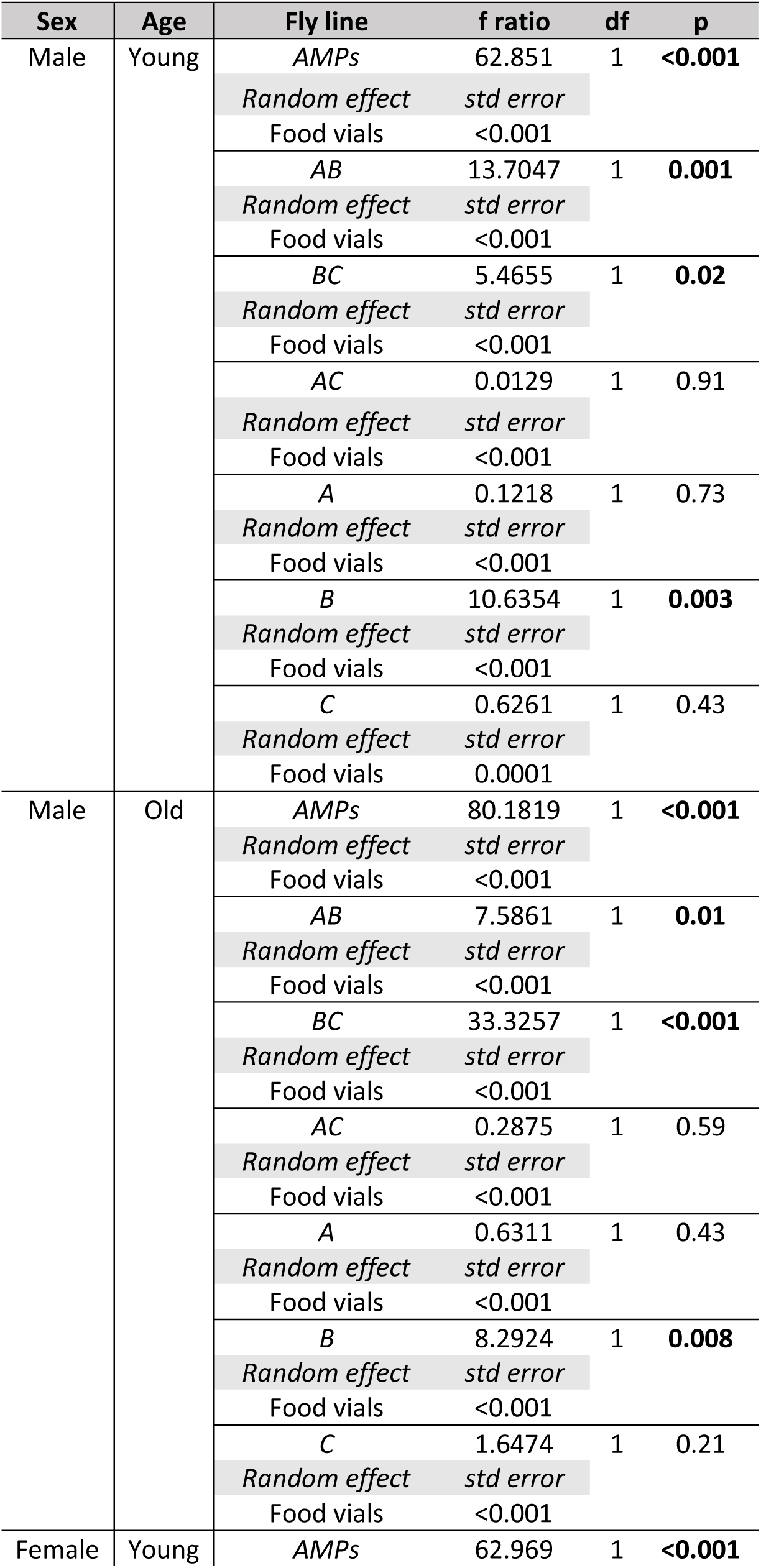
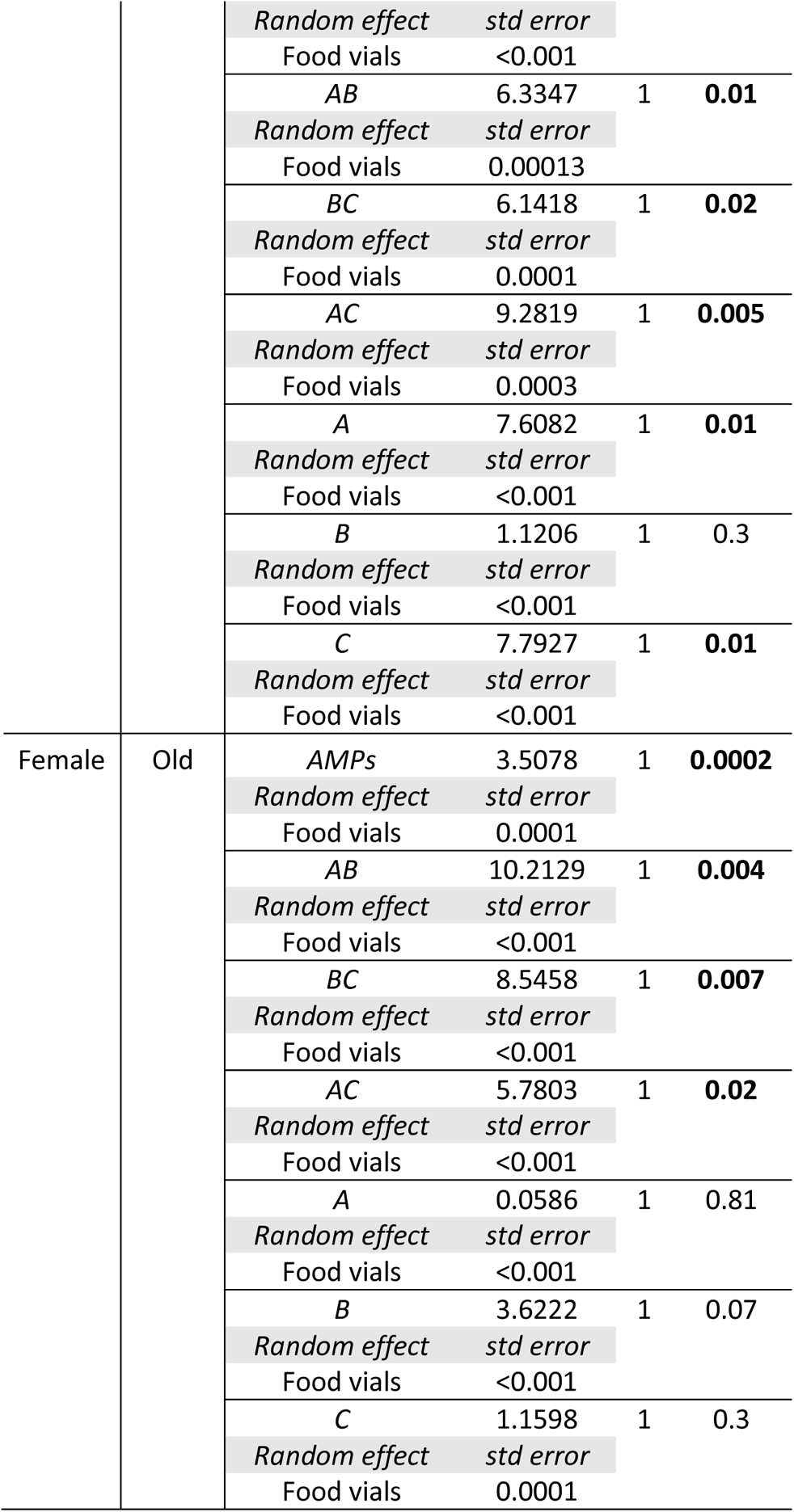
Summary of a generalized linear mixed model on log transformed bacterial load data (confirmed and fitted with a gamma distribution) for individual fly lines with compound mutations (i.e., A, B, C, AB, BC & AC) across sexes and age-groups, relative to control iso-*w^1118^* flies after *P. entomophila* infection. We specified the model as: log transformed bacterial load ~ fly line (individual mutant lines vs iso-*w^1118^*), with ‘fly line’ as fixed and ‘replicate food vials’ as random effects across sexes and age-groups separately.

**Table S9.**
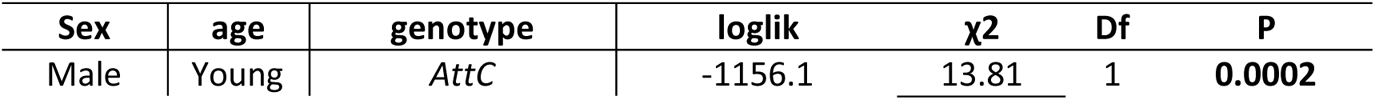
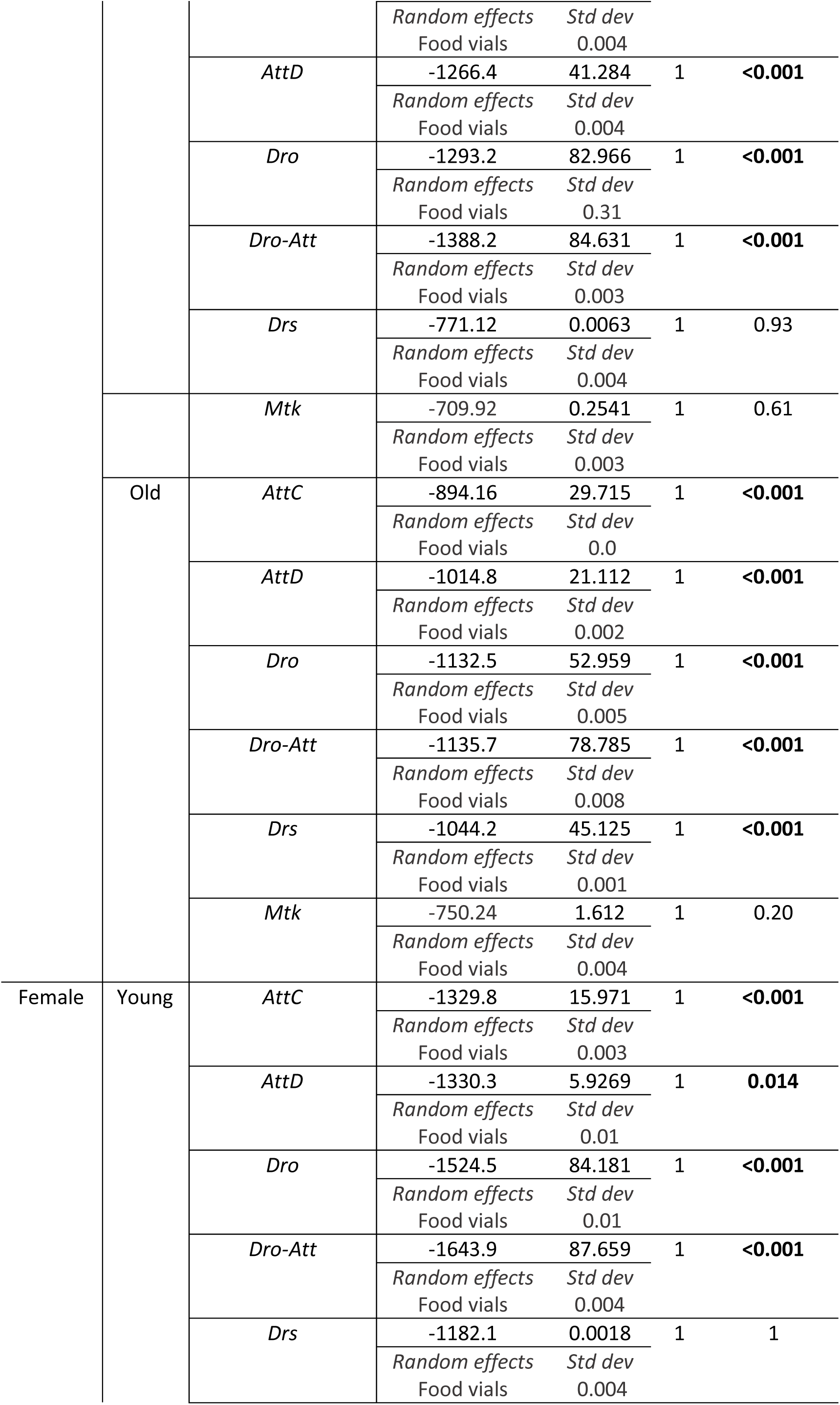
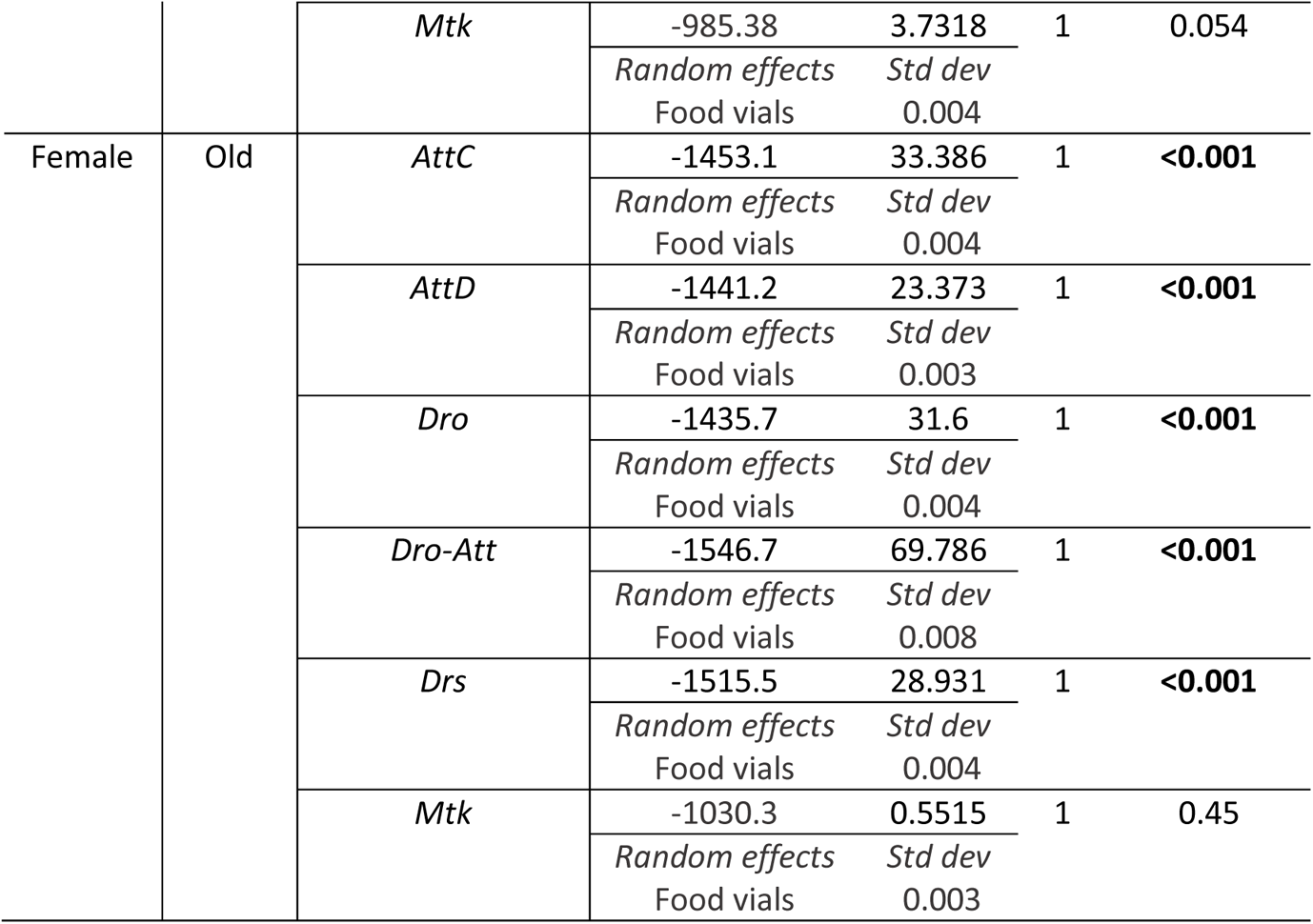
Summary of the mixed effects Cox model analyses to estimate the changes in the survival of individual Imd- and Toll-pathway specific and *Dro-Att* mutations across sexes and age-groups, relative to control iso-*w^1118^*flies after *P. entomophila* infection. For each fly line across sexes and age-groups, we specified the model as: survival ~ fly line (individual fly lines vs iso-*w^1118^*) + (1|food vials), with ‘fly line’ as a fixed effect and ‘replicate food vials’ as a random effect.

**Table S10.**
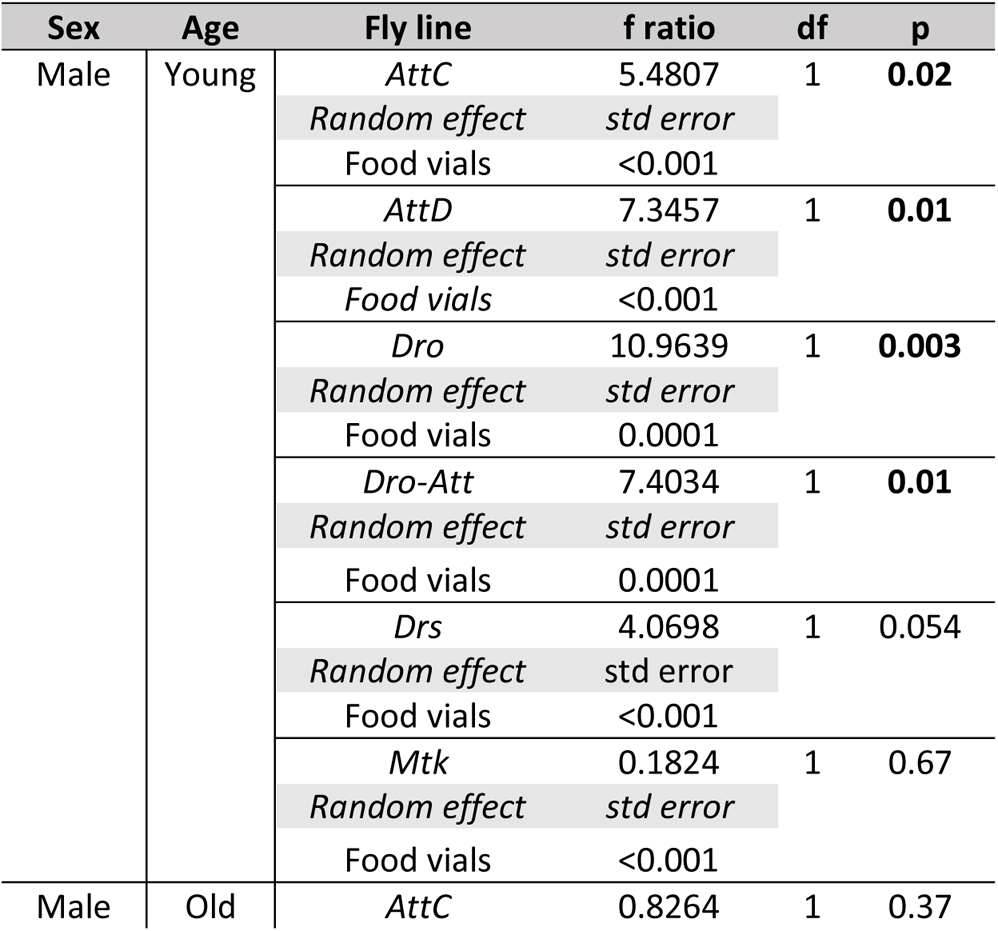
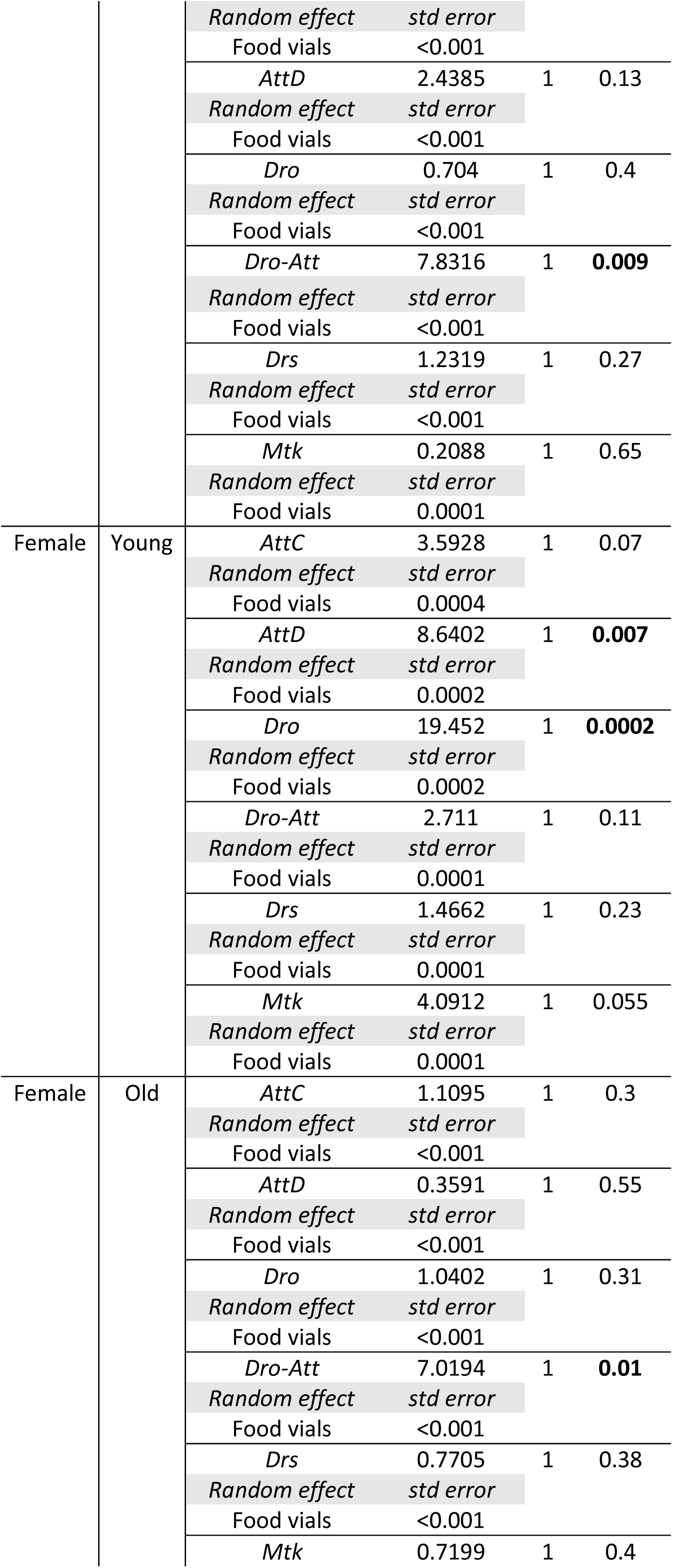
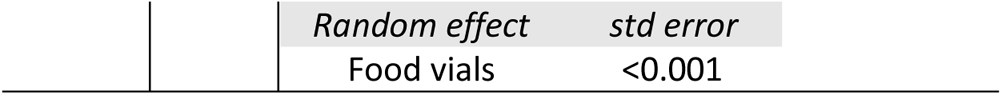
Summary of a generalized linear mixed model on log transformed bacterial load data (confirmed and fitted with a gamma distribution) for individual fly lines with Imd- & Toll-pathway specific, and *Dro-Att* mutations across sexes and age-groups, relative to control iso-*w^1118^* flies after *P. entomophila* infection. We specified the model as: log transformed bacterial load ~ fly line (individual mutant lines vs iso-*w^1118^*), with ‘fly line’ as fixed and ‘replicate food vials’ as random effects across sexes and age-groups separately.

**Table S11.** Summary of a generalized linear model (best fitted to a quasi-binomial distribution) for Malpighian tubule activity as a function of infection status across age-groups. For each age group, we specified the model as Malpighian tubule activity~ Infection status, with infection status as a fixed effect across age-groups.

| Age | Df | $\chi^2$ | P |
| --- | --- | --- | --- |
| Young | 1 | 0.0814 | 0.77 |
| Old | 1 | 6.179 | **0.012** |

**Table S12.**
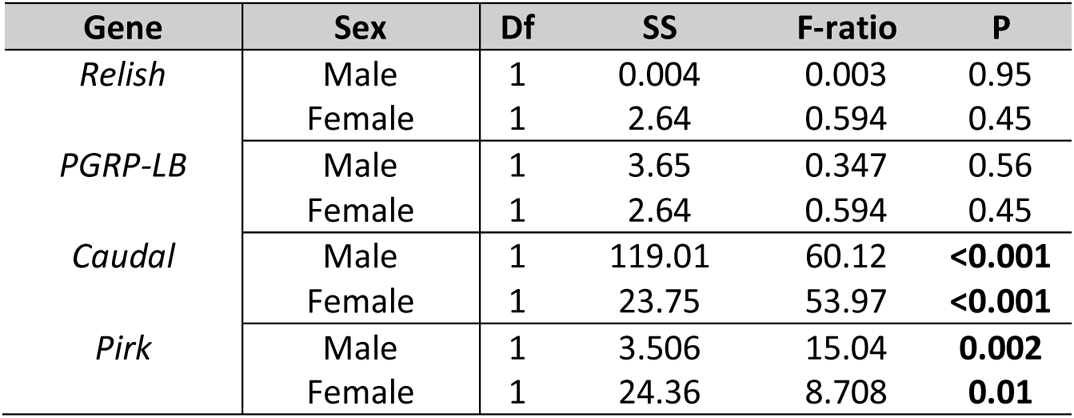
Summary of ANOVA on the gene expression data in males and females. For each sex and genes, we specified the model as: Fold-change in gene expression ~ age, with ‘age’ as a fixed effect.

